# Effects of alcohol on the transcriptome, methylome, and metabolome of *in vitro* gastrulating human embryonic cells

**DOI:** 10.1101/2024.09.26.615159

**Authors:** E Wallén, K Rämö, J Vehviläinen, J Sokka, M Lehtonen, T Otonkoski, R Trokovic, P Auvinen, O Kärkkäinen, N Kaminen-Ahola

**Affiliations:** Environmental Epigenetics Laboratory, Department of Medical and Clinical Genetics, Medicum, Faculty of Medicine, University of Helsinki, Helsinki, Finland; Research Programs Unit, Stem Cells and Metabolism and Biomedicum Stem Cell Centre, Faculty of Medicine, University of Helsinki, Helsinki, Finland; School of Pharmacy, Faculty of Health Sciences, University of Eastern Finland, Kuopio, Finland; Children’s Hospital, Helsinki University Central Hospital, University of Helsinki, Helsinki, Finland

**Keywords:** prenatal alcohol exposure, FASD, gastrulation, DNA methylation, embryonic development, mesoderm, endoderm, ectoderm

## Abstract

Prenatal alcohol exposure (PAE) affects embryonic development, causing a variable fetal alcohol spectrum disorder (FASD) phenotype with neurodevelopmental disorders and birth defects. To explore the effects of PAE on gastrulation, we used an *in vitro* model with subchronic moderate (20 mM) and severe (70 mM) ethanol exposures during the differentiation of human embryonic stem cells into germ layer cells. We analysed genome-wide gene expression (mRNA sequencing), DNA methylation (EPIC Illumina microarrays), and metabolome (non-targeted LC-MS method) of the endodermal, mesodermal, and ectodermal cells. The largest number of ethanol-induced alterations were observed in the endodermal cells, whereas the most prominent changes were seen in the ectodermal cells. Genes of the major morphogen signaling pathways involved in gastrulation and body patterning were affected by ethanol. Many of the altered genes, such as *BMP4*, *FGF8*, *SIX3,* and *LHX2*, have been previously associated with PAE and phenotypes of FASD, like defects in heart and corpus callosum development as well as holoprosencephaly. Furthermore, methionine metabolism was altered in all germ layer cells. Our findings support the early origin of alcohol-induced developmental disorders and strengthen the role of methionine cycle in the etiology of FASD.

## INTRODUCTION

Prenatal alcohol exposure (PAE) is one of the most significant causes of developmental disability affecting 3–5% of individuals in the Western world (<u>Roozen et al. 2016</u>). PAE can cause a wide range of developmental defects including neurodevelopmental disorders (NDDs), structural malformations, and growth disturbances referred to as Fetal Alcohol Spectrum Disorders (FASD). Its most severe form is fetal alcohol syndrome, which is characterized by craniofacial malformations, central nervous system defects as well as prenatal and postnatal growth restriction (<u>Hoyme et al. 2016</u>).

Many teratogenic effects of alcohol on the developing embryo have been traceable back to gastrulation (<u>Goodlett & Horn 2001</u>; <u>Lipinski et al., 2012</u>), resulting in a high incidence of FASD (<u>Sulik 1984</u>; <u>Maier & West 2001</u>; <u>Guerri 2002</u>). Gastrulation occurs in the third week after fertilization in human, and it is one of the most critical events in development, establishing directionality within the developing embryo and priming the system for organogenesis. During gastrulation, embryonic stem cells differentiate into three embryonic germ layers, of which endoderm, the innermost layer of the embryo, forms the linings of the gastrointestinal, respiratory, and urinary tracts. The middle layer, the mesoderm, forms the connective tissue, smooth muscle, cardiovascular system, skeleton, blood, and reproductive systems. The outermost layer is ectoderm, from which develops the external ectoderm, as well as the neuroectoderm including spinal cord, neural crest, and neural tube-brain. Alcohol-induced changes in the ectodermal cells are of particular interest, as the developing nervous system is known to be especially sensitive to the effects of PAE (<u>Guerri 2002</u>) and alcohol-related NDDs appear to be the most common diagnosis among the FASD (<u>Wozniak et</u> <u>al., 2019</u>). However, the effects of alcohol exposure on the embryonic germ layers in human are mostly unknown, as technical and ethical limitations challenge the study of this period.

The spatiotemporal control of gene expression is essential for directing cell type specification, migration, and localization. DNA methylation (DNAm) is a covalent epigenetic modification on the DNA strand, which by affecting chromatin density and the binding of transcription factors regulates gene expression according to the cell type and developmental stage. Early pregnancy, a period of dynamic cell divisions, DNA replication, and establishment of cell type-specific epigenetic profiles during epigenetic reprogramming, is particularly sensitive to environmentally induced epigenetic alterations (<u>Susiarjo et al., 2013</u>; <u>Tobi et al., 2015</u>). The establishment of DNAm profiles and their maintenance in mitotic cell divisions are accomplished by DNA methyltransferase enzymes. Early alcohol-induced DNAm alterations have been observed in human and mouse embryonic stem cells (hESCs and mESCs, respectively) (<u>Sánchez-Alvarez et al., 2013</u>; <u>Khalid et al. 2014</u>; <u>Auvinen et al., 2022</u>) as well as in the offspring of the early PAE mouse model (<u>Kaminen-Ahola et al., 2010</u>; <u>Zhang</u> <u>et al., 2015,</u> <u>Bestry et al., 2024</u>). It has been hypothesised that in addition to immediate cell death or DNA strand breaks, PAE can have an impact on the cells’ methylation potential and consequently alters the sensitive period of epigenetic reprogramming in early pregnancy. Previous studies have suggested that alcohol can affect methylation capacity through methionine metabolism (<u>Kruman et</u> <u>al., 2014</u>; <u>Ngai et al., 2015</u>), and by interfering with the activity of DNA methyltransferases (<u>Bönsch,</u> <u>et al., 2006</u>; <u>Miozzo et al., 2018</u>).

Here we elucidate the effects of ethanol (EtOH) exposure on early development by exposing hESCs to moderate (20 mM) and severe (70 mM) subchronic EtOH exposure while differentiating them into the endodermal, mesodermal, and ectodermal cells. We explored the effects of EtOH on the transcriptome, methylome, and metabolome of the embryonic germ layer cells.

## RESULTS

### Effects of EtOH on the transcriptome of germ layer cells

To study genome-wide EtOH-induced alterations in gene expression, we performed 3’mRNA-sequencing (mRNA-seq) for controls as well as 20 mM and 70 mM EtOH-exposed cells of three germ layers (*n* = 4 replicates/germ layer). When cells were exposed to 20 mM EtOH, differentially expressed genes (DEGs) (FDR < 0.05) were observed only in endodermal cells (79 DEGs of which 69 were down– and 10 upregulated) (Table S1).

#### Endodermal cells

The largest number of significant changes in the germ layers were observed in the 70 mM EtOH-exposed endodermal cells including a total of 154 DEGs (67 down– and 87 upregulated) of which 30 were common with the cells exposed to 20 mM EtOH (Fig. 1a, Table S1). When 70 mM EtOH-exposed cells were compared to controls, a highly upregulated *Nodal Growth Differentiation Factor* (*NODAL*), *Cerberus 1* (*CER1*) as well as *Left-Right Determination Factors 1* and *2* (*LEFTY1* and *LEFTY2*) were observed. They all are involved in the TGF-β signaling pathway as well as the left-right axis determination in a developing zebrafish embryo (<u>Smith et al. 2011,</u> <u>Hashimoto et al., 2004</u>). This process is crucial in establishing correct patterning for organ development, and interestingly, the loss of left-right symmetry as well as midline defects have been associated with PAE (<u>Klingenberg et al., 2010</u>). Also, we observed a highly upregulated *Insulin Like Growth Factor Binding Protein 5* (*IGFBP5*), which regulates the growth and development of cells and tissues in the early mouse embryos (<u>Salih et al., 2004</u>).

**Fig. 1.**
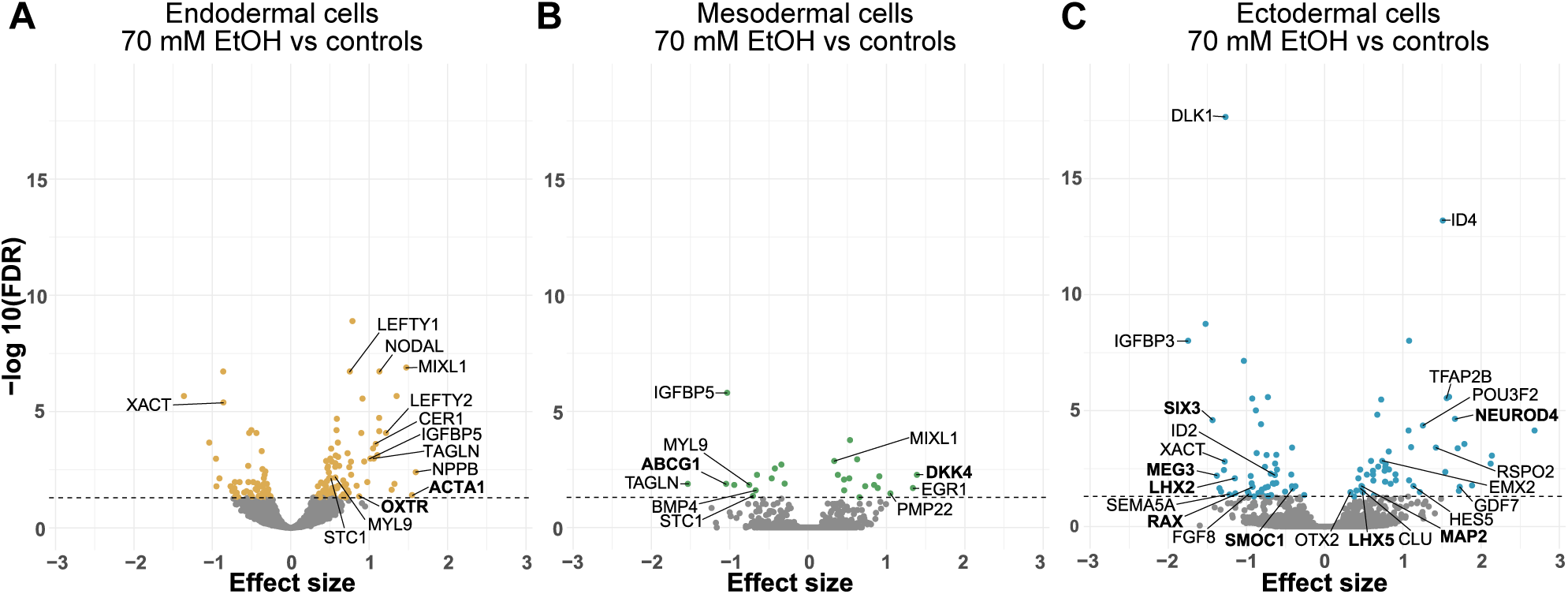
Differential gene expression in the 70 mM EtOH-exposed germ layer cells. **a-c** Volcano plots showing the effect of 70 mM EtOH exposure on mRNA expression in the **a** endodermal cells, **b** mesodermal cells and **c** ectodermal cells. Horizontal line marks FDR 0.05. Common significantly altered genes between DNAm and mRNA-seq analyses in each germ layer are bolded.

To get an overall picture of the processes in which EtOH-induced DEGs cluster, we performed Gene Ontology (GO) pathway enrichment analysis for biological process (BP), molecular function (MF), and cellular component (CC). The most significant BP term for both EtOH exposures was a ribosome-mediated process ‘cytoplasmic translation’ (FDR-corrected q-value < 0.05, Table S2). In the 70 mM EtOH-exposed endodermal cells, the cholesterol metabolic process, sterol metabolic process, regulation of microtubule polymerization, and cell migration involved in gastrulation as well as alcohol metabolic process BP terms were also significant (Fig. 2a).

**Fig. 2.**
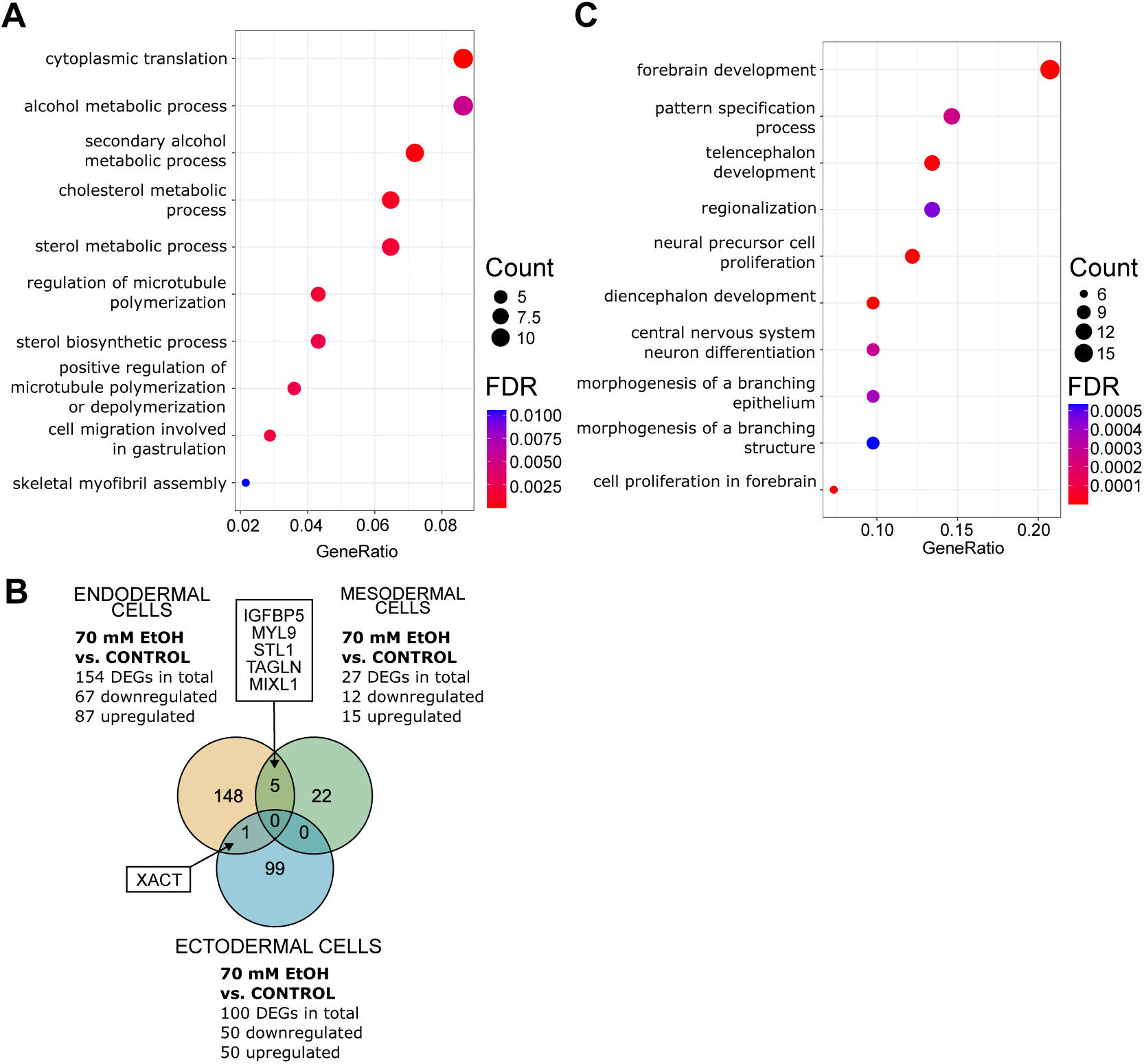
Gene expression pathways and common genes of the 70 mM EtOH-exposed germ layer cells. **a** Significantly enriched terms identified in GO:BP enrichment analysis of 70 mM EtOH-induced DEGs in endodermal cells (FDR-corrected q-value < 0.05). The 10 most significant pathways are shown. **b** Venn diagram showing the number of 70 mM EtOH-induced DEGs, which are in common between germ layers. **c** Significantly enriched terms identified in GO:BP enrichment analysis of 70 mM EtOH-induced DEGs in ectodermal cells (FDR-corrected q-value < 0.05). The 10 most significant pathways are shown.

#### Mesodermal cells

The lowest number of significant changes were observed in the mesodermal cells. We detected 27 DEGs (12 down– and 15 upregulated), including downregulated *BMP4, IGFBP5*, *TAGLN*, and *ABCG1* as well as upregulated *DKK4* and *EGR1* in the 70 mM EtOH-exposed cells (Fig. 1b, Table S3). *Bone morphogenetic protein 4 (BMP4)* is a key ligand that drives the mesoderm formation and an inducer of early cardiac mesoderm (<u>Tsaytler et al. 2023</u>). Previously, reduced Bmp signaling has been observed in the heart cone and heart tube of EtOH-exposed zebrafish embryos (<u>Sarmah et al. 2016</u>). Furthermore, *ATP-binding cassette sub-family G member 1* (*ABCG1*) has been linked to PAE in human and rat placenta (<u>Auvinen et al., 2022,</u> <u>Subbanna & Basavarajappa et al.,</u> <u>2022</u>) and *Early Growth Response 1* (*EGR1*) in mouse embryonic forebrain (<u>Zhang et al., 2019)</u>. Interestingly, even though there were not many DEGs in the mesodermal cells, five of them, *IGFBP5*, *MIXL1*, *MYL9*, *STC1*, and *TAGLN,* were in common with the endodermal cells (Fig. 2b). One of these, upregulated *Mix Paired-Like Homeobox* (*MIXL1*), is a transcription factor expressed in the primitive streak of the gastrulating embryo, and it plays a critical role in marking the cells to form mesoderm and endoderm (<u>Ng et al., 2005</u>).

#### Ectodermal cells

Based on effect sizes, the most prominent changes in gene expression were observed in the 70 mM EtOH-exposed ectodermal cells with 100 DEGs (50 down– and 50 upregulated) (Fig. 1c, Table S4). Among the DEGs were homeobox genes *EMX2*, *LHX2*, *LHX5*, *OTX2*, *POU3F2*, *RAX*, and *SIX3*, growth associated factors *GDF7, IGFBP3,* and *FGF8,* as well as other developmentally essential genes such as *DLK1*, *MEG3*, *SEMA5A*, *NEUROD4, ID2* and *ID4*. *Orthodenticle homeobox 2* (*OTX2),* a transcription factor involved in the formation of neuroectoderm during gastrulation (<u>Rhinn et al., 1998</u>; <u>Larsen et al., 2010</u>) and craniofacial development (<u>Matsuo et</u> <u>al., 1995</u>), has been previously associated with PAE in mouse brain (<u>Kleiber et al. 2016</u>). The expression of *Six homeobox 3* (*SIX3*), a transcription factor essential for the forebrain formation as well as the expressions of *EMX2, LHX2, LHX5,* and *OTX2,* were all altered by EtOH in *in vitro* human pluripotent stem cell-based model of corticogenesis (<u>Fisher et al. 2021</u>). *Fibroblast growth factor 8* (*FGF8)* belongs to FGF signaling pathway, which is crucial for various tissues and organ systems, including the development of craniofacial features, cardiovascular structures, and brain development (<u>Xie et al., 2020</u>) commonly affected in FASD (<u>Hoyme et al., 2016</u>). Also, previous human and mouse PAE studies have found alterations in the same genes as in the current work, including *EMX2* (<u>Rosenberg et al., 2010</u>), *LHX2* (<u>Subbanna & Basavarajappa et al., 2022</u>), *ID2* (<u>Abbott et al., 2018</u>; <u>Bottom et al., 2022</u>; <u>Conner et al., 2020</u>; <u>El Shawa et al., 2013;</u> <u>Hashimoto-Torii et al., 2011</u>), *ID4* (<u>Rosenberg et al., 2010;</u> <u>Boschen et al., 2021</u>), *RSPO2* (<u>Sambo et al., 2022</u>), *MAP2* (<u>Putzke et al.,</u> <u>1998</u>), and *CLU* (<u>Kim et al., 2012</u>).

The most significantly downregulated paternally expressed *delta like non-canonical Notch ligand 1 (DLK1)* as well as *maternally expressed 3* (*MEG3*) locate at the *DLK1-DIO3* locus. They both belong to imprinted genes, which are epigenetically regulated according to the parent of origin in a locus-specific manner. This locus is necessary for fetal development and postnatal growth, and its ncRNAs are highly expressed in early embryo (<u>Mo et al., 2015</u>; <u>Prats-Puig et al., 2017</u>). There was one common DEG, downregulated *X Active Specific Transcript* (*XACT*), between 70 mM EtOH-exposed endodermal and ectodermal cells (Fig 2b). *XACT* is a long non-coding RNA, which plays a role in the control of X-chromosome inactivation (<u>Vallot et al., 2013</u>).

Ectodermal GO:BP pathways are closely related to the brain, nervous system, and eye development, the most significant being forebrain development, cell proliferation in forebrain, neural precursor cell proliferation, diencephalon development, telencephalon development, pattern specification process, and central nervous system neuron differentiation (q-value < 0.05, Table S5, Fig. 2c).

### Effects of EtOH on the methylome of germ layer cells

We used DNAm microarrays (Infinium MethylationEPIC BeadChip, Illumina) to explore the effects of EtOH exposure on the genome-wide DNAm of germ layer cells (control *n* = 4 replicates/germ layer, 70 mM EtOH *n* = 4 replicates/germ layer). Since the transcriptome of all germ layers was significantly altered only at 70 mM exposure, we focused only on this concentration in the methylome analyses. We calculated genome-wide average DNAm (GWAM) by using all probes in the array and did not observe significant EtOH-induced alterations in total DNAm levels (Fig. S1). However, in the mesodermal cells, we observed hypomethylation in relation to islands: north shores (N_shore) and south shores (S_shore) (*P* = 0.044 and *P* = 0.041, respectively, Student’s *t* test, Fig. S1b), and in the ectodermal cells, we observed significant hypomethylation at genomic location 200 bp upstream of transcription start site (TSS200) (*P* = 0.046, Welch’s Two-Sample *t* test, Fig. S1c). The global DNAm level was also predicted by comparing the mean DNAm level of CpGs in Alu, endogenous retrovirus (ERV), and long interspersed nuclear element 1 (LINE1) repetitive elements (RE), which comprise 32% of human genome in total (<u>Zheng et al., 2017</u>; <u>International Human Genome</u> <u>Sequencing Consortium, 2001</u>). When EtOH-exposed ectodermal cells were compared to controls, significantly hypermethylated ERV regions were observed (*P* = 0.035, Student’s *t* test) (Fig. S2). This is in line with our previous results where long terminal repeats (LTRs), which also include ERVs, were hypermethylated in a mouse model of early alcohol exposure (<u>Kaminen-Ahola et al., 2010</u>) as well as in EtOH-exposed hESCs and human PAE placenta (<u>Auvinen et al., 2022</u>).

#### Endodermal cells

Similarly to gene expression analyses, the largest number of significant changes in DNAm analyses were observed in the endodermal cells with 568 EtOH-induced differentially methylated CpG sites (hereafter differentially methylated positions, DMPs, with FDR < 0.05) (131 hypo– and 437 hypermethylated, associated with a total of 93 and 341 genes, respectively) (Fig. 3a, Table S6). Several homeobox genes *ALX4*, *LHX3*, *LHX5*, *NKX2*-5, *HOXB13*, *PAX5,* and *PAX7* as well as WNT gene family members *WNT3A*, *WNT6,* and *WNT7B,* which all play crucial roles for many developmental processes in gastrulation, were hypermethylated in endodermal cells. Also, genes encoding zinc finger proteins, which function in transcriptional regulation during early embryonic development and cell differentiation (<u>Cassandri et al., 2017</u>), were altered in endodermal cells, including hypomethylated *ZNF274*, *ZNF454*, *ZNF502*, *ZNF528*, *ZNF578*, *PRDM7*, and hypermethylated *PRDM16*. *PRDM7* and *PRDM16* are both histone methyltransferases and critical epigenetic regulators of early development (<u>Pinheiro et al., 2012</u>; <u>Di Tullio et al., 2022</u>). Owing to previous mouse and zebrafish studies, *PRDM16* plays a major role in craniofacial development (<u>Shull</u> <u>et al., 2020</u>), abnormalities of which are characteristic of FASD (<u>Hoyme et al. 2016</u>). Several altered genes have been associated previously with EtOH exposure or PAE, such as *ALX4* in embryonic mouse heart (<u>Abraham et al., 2022</u>), *SIM1* at early neurulation in mouse embryo (<u>Liu et al., 2009</u>) as well as *FOXP2* and genes in *WNT* signaling pathway in human PAE placenta in our previous study (<u>Auvinen et al., 2022</u>).

**Fig. 3.**
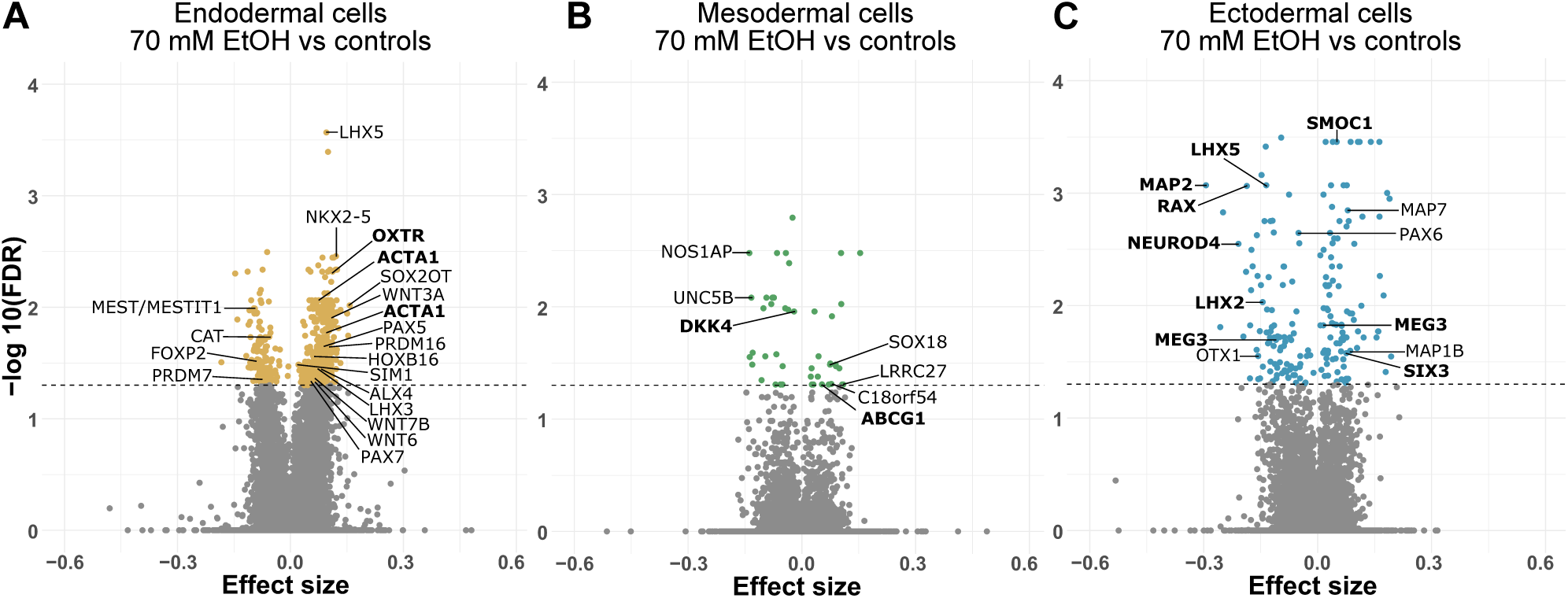
Differential DNAm of the 70 mM EtOH-exposed germ layer cells. **a-c** Volcano plots showing the effect of 70 mM EtOH exposure on DNAm in the **a** endodermal cells, **b** mesodermal cells, and **c** ectodermal cells. Horizontal line marks FDR 0.05. Common significantly altered genes between DNAm and mRNA-seq analyses in each germ layer are bolded.

Moreover, 56 differentially methylated regions (DMRs), defined as a region with maximal allowed genomic distance of 1,000 bp containing three or more CpGs, were observed in endodermal cells (Table S7). The most significantly hypomethylated DMR as well as seven hypomethylated DMPs in regulatory and promoter regions associated with imprinted *Mesoderm Specific Transcript* (*MEST*, also known as *PEG1*) and its antisense RNA *MEST Intronic Transcript 1* (*MESTIT1*), which shares a promoter region with *MEST* (<u>Nakabayashi et al., 2002</u>). It is paternally expressed during embryonic development, and it encodes a member of the alpha/beta hydrolase superfamily (<u>Kobayashi et al.,</u> <u>1997</u>). The most prominently hypermethylated DMP and two DMRs in the endodermal cells were in non-coding RNA *SOX2-OT*. It regulates the expression of *SOX2*, which is a well-known developmentally critical transcription factor for maintaining pluripotency and directing differentiation into the germ layers (<u>Masui et al., 2007</u>; <u>Zhang et al., 2019</u>). Interestingly, previous studies by mESCs (<u>Ogony et al., 2013</u>; <u>Sánchez-Alvarez et al., 2013</u>) and human placenta (<u>Auvinen</u> <u>et al., 2022</u>) suggest that EtOH can reprogram the lineage specification by changing the dosage of pluripotency factors Oct4 and Sox2. Furthermore, we detected a hypomethylated DMR in *CAT* and hypermethylated DMRs in *LHX5* and *NKX2-5*. Catalase, encoded by *CAT*, is an important antioxidant enzyme protecting against alcohol-induced damage and alleviating oxidative stress (<u>Brocardo et al.,</u> <u>2011</u>; <u>Miller et al., 2013</u>), and it is also associated with PAE in mouse brain and heart (<u>Drever et al.,</u> <u>2012</u>; <u>Atum et al., 2023</u>).

We performed GO pathway enrichment analysis for BP, MF, and CC, and the most significant terms were related to MFs, such as transcription factor activity and transcription regulatory region binding (*P* < 0.05, Table S8, Fig. 4a). Interestingly, there were also several significant terms related to neurodevelopment, such as spinal cord association neuron differentiation, nervous system development, generation of neurons, and neuron differentiation.

**Fig. 4.**
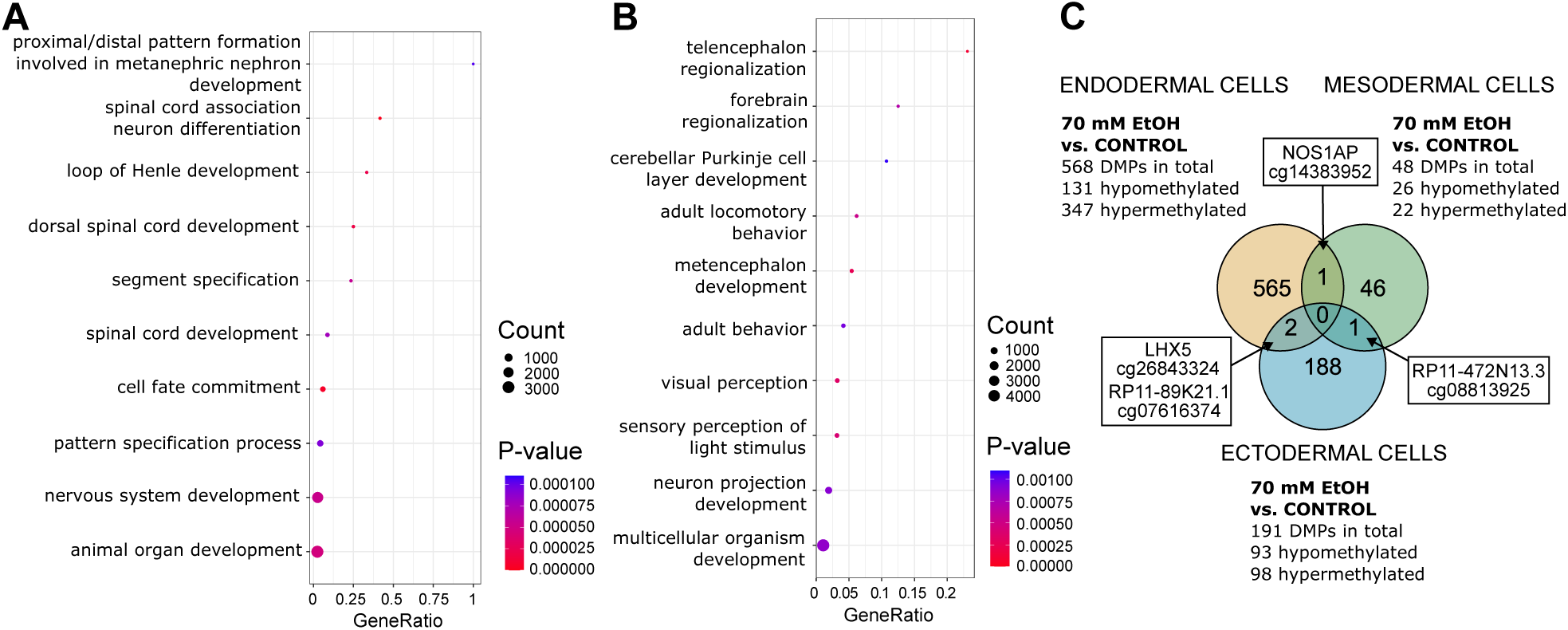
DNAm pathways and common DMPs of the 70 mM EtOH-exposed germ layer cells. **a** Significantly enriched terms identified in GO:BP enrichment analysis of 70 mM EtOH-induced DMPs in endodermal cells (*P* < 0.05). The 10 most significant pathways are shown. **b** Significantly enriched terms identified in GO:BP enrichment analysis of 70 mM EtOH-induced DMPs in ectodermal cells (*P* < 0.05). The 10 most significant pathways are shown. **c** Venn diagram showing the number of 70 mM EtOH-induced DMPs, which are in common between germ layers.

#### Mesodermal cells

In line with the low number of alterations in mRNA-seq analysis, we observed only 48 EtOH-induced DMPs (26 hypo– and 22 hypermethylated associated with 14 and 15 genes, respectively) (Fig. 3b, Table S9) in the mesodermal cells. Additionally, six DMRs associated with *TOLLIP*, *ASB18*, *CHRND*, and *DDR2* were observed (Table S10). The most prominently hypomethylated DMP was in *Nitric Oxide Synthase 1 Adaptor Protein* (*NOS1AP*), which encodes a protein involved in neurotransmitter nitric-oxide synthesis (NOS) together with *NOS1*. The reduction of NOS activity in developing brain in PAE rodent models has been reported previously (<u>Kimura et</u> <u>al., 1996</u>; <u>Kimura & Brien, 1998</u>; <u>Bonthius et al., 2002</u>; <u>de Licona et al., 2009</u>). Moreover, *NOS1AP* was hypermethylated in buccal epithelial cells in children with FASD (<u>Portales-Casamar et al., 2016</u>). *NOS1AP* together with the *NOS1* are also associated with several psychiatric disorders, such as attention-deficit/hyperactivity disorder (ADHD) (<u>Freudenberg et al., 2015</u>). Several other altered genes in the mesodermal cells are also associated with EtOH exposure in previous studies, such as *UNC5B, C18orf54,* and *SOX18* in the gastrulation-stage mouse embryo (<u>Boschen et al., 2021</u>), *SOX18* in hESCs (<u>Sánchez-Alvarez et al., 2013</u>) as well as *LRRC27* and *DDR2* in human dental pulp stem cells (<u>Hoang et al., 2016</u>).

#### Ectodermal cells

We observed 191 EtOH-induced DMPs (93 hypo– and 98 hypermethylated associated with 70 and 67 genes, respectively, Fig. 3c, Table S11) and 28 DMRs in the ectodermal cells (Table S12). Similarly to DEGs in mRNA-seq analyses, there were fewer DMPs in the ectodermal cells compared to endodermal cells, but the ectodermal DMPs were more prominently altered based on effect sizes. The most prominently hypomethylated DMP was in the 5’ untranslated region (5’UTR) of *Microtubule Associated Protein 2* (*MAP2*). This gene encodes a neuron-specific cytoskeletal protein that plays an important role in stabilizing dendritic shape and is a known marker of dendrites and neuronal health (<u>Dehmelt et al., 2005</u>; <u>DeGiosio et al., 2022)</u>. *MAP2* has been associated with several neuropsychiatric and neurodegenerative disorders (<u>DeGiosio et al., 2022</u>) as well as alterations in EtOH-exposed rat brain and hippocampal cultured neurons (<u>Putzke et al., 1998</u>; <u>Romero et al., 2010</u>), and PAE rat frontal cortex (<u>Tan et al., 1993</u>). Additionally, two other microtubule-associated genes, *MAP1B* and *MAP7*, were both hypermethylated in the ectodermal cells.

Furthermore, there were many homeobox genes altered by EtOH, including hypomethylated *LHX2*, *LHX5*, *PAX6*, *OTX1*, and *RAX* as well as hypermethylated *SIX3*. A transcription factor *LIM homeobox 2 (LHX2)* regulates the neural differentiation of hESCs via transcriptional modulation of *Paired box 6* (*PAX6)* and *CER1* (<u>Hou et al. 2013</u>). *PAX6* has been associated with EtOH exposure or PAE in numerous studies (<u>Mo et al., 2012</u>; <u>Fisher et al., 2021; Kim et al., 2010</u>; <u>Veazey et al., 2013; Aronne</u> <u>et al., 2008;</u> <u>Zhang et al., 2014</u>; <u>Peng et al., 2004</u>; <u>Yelin et al., 2007</u>), *LHX2* and *PAX6* have been associated previously with PAE in mouse forebrain (<u>Subbanna & Basavarajappa et al., 2022</u>), and *OTX1* with PAE in the embryonic mouse brain (<u>Zheng et al., 2014</u>).

GO pathway enrichment analysis of EtOH-induced DMPs revealed expectedly, that almost all terms in ectodermal cells were related to brain and nervous system development such as telencephalon regionalization and metencephalon development, visual perception as well as forebrain regionalization. Interestingly, also BP terms adult locomotory behaviour and adult behaviour were seen (Table S13, Fig. 4b).

Moreover, we compared EtOH-induced alterations between the cells of three germ layers. There was one common DMP between endodermal and mesodermal cells, *NOS1AP* (cg14383952), which was hypomethylated in both germ layers (Fig. 4c). Also, one common DMP was observed in mesodermal and ectodermal cells associated with intergenic long non-coding RNA *RP11-472N13.3* (cg08813925), which was hypermethylated in both germ layers. Furthermore, there were two common DMPs in endodermal and ectodermal cells: *LHX5* (cg26843324) and *RP11-89K21.1* (cg07616374) were hypermethylated in the endodermal and hypomethylated in the ectodermal cells.

Finally, gene expression results were compared with DNAm results, and in the endodermal cells, there were two common genes: *ACTA1* (five DMPs in TSS200 and TSS1500) and *OXTR* (DMP in 5’UTR) that were hypermethylated and upregulated. Mesodermal cells share two common genes between analyses, hypomethylated and upregulated *DKK4* (DMP in TSS1500), and hypermethylated and downregulated *ABCG1* (DMP in 5’UTR). There were eight genes in common in the ectodermal cells, of which three were hypomethylated and upregulated: *LHX5* (two DMPs in gene bodies), *MAP2* (DMP in 5’UTR), and *NEUROD4* (DMP in 5’UTR), two hypermethylated and downregulated: *SIX3* (DMP in gene body) and *SMOC1* (DMP in gene body), and two hypomethylated and downregulated: *LHX2* (three DMPs in TSS200, TSS1500 and 5’UTR) and *RAX* (two DMPs in 3’UTR). Two hypomethylated and one hypermethylated DMPs of *MEG3* (DMPs in 5’UTR) and downregulated gene expression were observed.

### Effects of EtOH on the metabolomics of germ layer cells

To elucidate the effects of EtOH on the metabolome and the interplay between methylome, transcriptome, and metabolome, we explored extracellular metabolites in the supernatants of the same germ layer cells (endodermal and mesodermal supernatants *n =* 4 replicates/condition and ectodermal supernatant *n* = 5 replicates/condition). We used a non-targeted LC-MS metabolomics method (<u>Klåvus et al., 2020</u>) to measure a total of 9,401 extracellular molecular features from germ layers. The altered metabolic profiles were unique in each germ layer (Fig. S3), and a clear EtOH dose response between EtOH concentrations was observed. Similar trends of alterations were seen in the transcriptome, methylome, and metabolome of the germ layer cells, as the largest quantitative changes were observed in the endodermal supernatant. Characteristics, statistical results, and reference spectra for all molecular features and identified metabolites are given as supplementary material (Table S14). Metabolite identification was focused on molecular features with *P* < 0.05 in comparisons between controls and EtOH exposures. Identified metabolites were classified into identification levels 1 and 2 according to Sumner et al. (<u>2007</u>). A total of 130 identified and significantly altered metabolites are shown in Table S15.

#### Endodermal supernatants

In the 20 mM EtOH-exposed endodermal cells, 62 metabolites with *P* < 0.05 (Welch’s *t* test) and four metabolites with FDR < 0.05 were observed, whereas in the 70 mM exposure, we observed 86 metabolites with *P* < 0.05 (Welch’s *t* test) and 32 metabolites with FDR < 0.05 (Fig. 5a). Compared to mesoderm and ectoderm, the levels of the altered metabolites in the endodermal supernatants were highly decreased, and the changes were visible especially in lysophosphocholines (LPCs) and lysophosphoethanolamines (LPEs). Almost all identified amino acids, including amino acid derivative glycine betaine as well as nucleosides were decreased, especially in the severe EtOH exposure.

**Figure 5.**
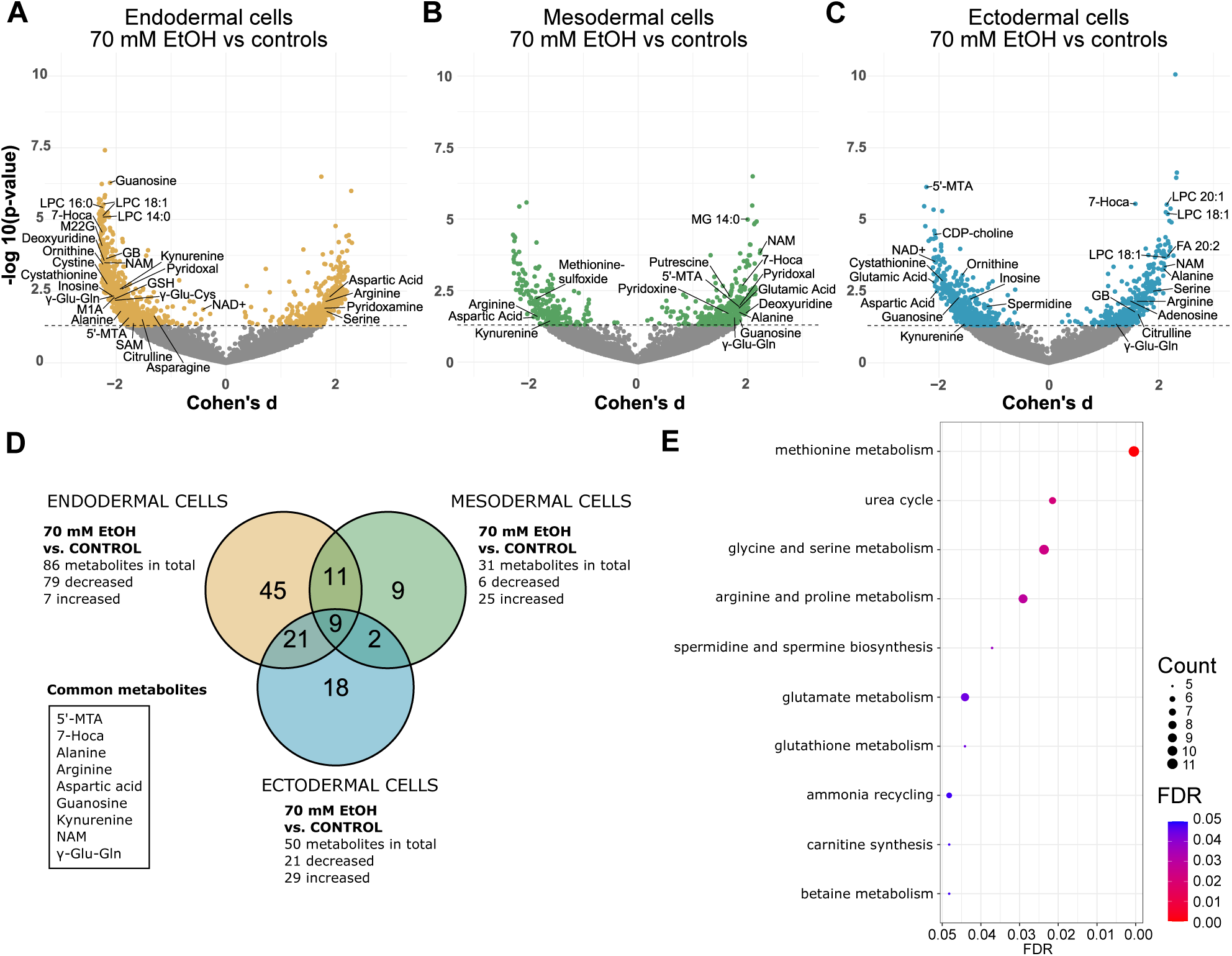
Altered metabolites in the 70 mM EtOH-exposed germ layers. **a-c** Volcano plots showing the effect of 70 mM EtOH exposure on extracellular metabolites in the **a** endodermal cells, **b** mesodermal cells and **c** ectodermal cells. Horizontal line marks *P*-value 0.05. **d** Venn diagrams showing the numbers of 70 mM EtOH-induced alterations in annotated metabolites. **e** Significantly enriched terms identified in SMPDB metabolite set enrichment analyses of all EtOH-induced annotated metabolites (*P* < 0.05). The 10 most significant pathways are shown (FDR < 0.05). 5’-MTA, 5’-methylthioadenosine; 7-Hoca, 7a-hydroxy-3-oxo-4-cholestenoic acid; GB, glycine betaine; GSH, reduced glutathione; LPC, lysophosphocholine; NAD+, nicotinamide adenine dinucleotide; NAM, N-acetylmethionine; M1A, 1-methyladenosine; M22G, N2,N2-dimethylguanosine; SAM, S-adenosylmethionine; γ-Glu-Cys, γ-glutamyl-cysteine; γ-Glu-Gln, γ-glutamyl-glutamine.

Furthermore, the amounts of S-Adenosylmethionine (SAM), a cofactor of DNA and histone methyltransferases, and nicotinamide adenine dinucleotide (NAD+), a redox reaction cofactor, were decreased in the endodermal supernatants (*P* < 0.05). SAM is a known principal methyl donor in numerous methylation reactions and cellular processes (<u>Lu, 2000</u>). Excessive alcohol consumption can decrease SAM levels, leading to aberrant DNAm patterns and alterations in gene expression (<u>Varela-Rey et al., 2013</u>). We also calculated the SAM/SAH ratio, commonly referred to as the methylation index, and observed a significantly decreased ratio in the severely exposed endodermal supernatants compared to controls (*P* = 0.036, Student’s *t* test, Fig. S4a). Observed significantly lowered asparagine levels in the 70 mM EtOH-exposed endodermal cells are in line with previous studies, where heavy alcohol consumption reduced the amount of asparagine in the human serum samples (<u>Kärkkäinen et al., 2020</u>). This is also observed in mothers who consumed alcohol during the first trimester of pregnancy (<u>Lehikoinen et al., 2018</u>). Interestingly, the increase in acetaldehyde due to alcohol consumption can induce decreased asparagine levels (<u>John et al., 2024</u>).

#### Mesodermal supernatants

In the 20 mM EtOH-exposed mesodermal cells, there were 20 metabolites with *P* < 0.05 (Welch’s *t* test), and in the 70 mM exposure, there were 31 metabolites with *P* < 0.05 (Welch’s *t* test) and only one metabolite, monoacylglyceride MG 14:0, with FDR < 0.05 (Fig. 5b). EtOH mostly increased metabolite levels in the 70 mM exposure, including two forms of vitamin B6, pyridoxal and pyridoxine, as well as polyamine putrescine. Changes in the diamine oxidase (DAO) activity, which is the rate-limiting enzyme in the terminal catabolism of polyamines, and the amounts of putrescine and other polyamines, such as spermidine, have been previously associated with alcohol consumption and PAE in rat (<u>Sessa & Perin, 1997</u>).

#### Ectodermal supernatants

Similarly to gene expression, the amounts of altered metabolites were more evenly increased or decreased in the ectodermal cells compared to the endodermal and mesodermal cells. In the supernatants of 20 mM EtOH-exposed ectodermal cells, 30 metabolites with *P* < 0.05 (Welch’s *t* test) were observed. In 70 mM exposure, we observed 50 metabolites with *P* < 0.05 (Welch’s *t* test) and eight metabolites with FDR < 0.05, which were decreased 5’-methylthioadenosine (5’-MTA), cytidine diphosphate choline (CDP-choline, also known as citicoline), and NAD+ as well as increased three LPCs, intermediate of bile acid synthesis 7a-hydroxy-3-oxo-4-cholestenoic acid, and fatty acid (FA) 20:2 (Fig. 5c).

N-acetylmethionine (NAM), a bioavailable source of methionine, was the only metabolite changed in all germ layers and EtOH concentrations in this study: it was significantly decreased in endodermal and significantly increased in mesodermal and ectodermal supernatants (*P* < 0.05). In addition to NAM, in the 20 mM exposure, there were two common altered metabolites between germ layers: deoxyuridine and hydroxyphenyllactate, and in the 70 mM exposure, there were eight common altered metabolites: 7a-hydroxy-3-oxo-4-cholestenoic acid, 5’-MTA, alanine, aspartic acid, arginine, γ-glutamylglutamine, guanosine and kynurenine (Fig. 5d). 5’-MTA, intermediate in the generation of adenine and methionine, and produced by the decarboxylation of SAM, was significantly less abundant in endodermal and ectodermal cells and more abundant in mesodermal cells compared to controls. 5’-MTA and kynurenine have neuroprotective effects in the human brain (<u>Moreno et al.,</u> <u>2014; Vamos et al., 2009</u>) and they have been associated with EtOH exposure in hESC-derived neural lineages (<u>Palmer et al., 2012</u>). Additionally, alterations in kynurenine have been recently associated with alcohol use disorder in several studies (<u>Vidal et al., 2020</u>, <u>Leclercq et al., 2021</u>, <u>Jang et al., 2022</u>, <u>Mechtcheriakov et al., 2022</u>).

We also observed several EtOH-induced alterations in the transsulfuration pathway, including cystathionine, cystine, γ-glutamyl-cysteine (γ-Glu-Cys), reduced glutathione (GSH) and oxidised glutathione (GSSG). GSH is a known protector against oxidative stress (<u>Hayes & McLellan, 1999</u>), and in a previous study, GSH level was elevated in EtOH-exposed neonatal rat brain (<u>Smith et al.,</u> <u>2005</u>). The GSH/GSSG ratio is an important indicator of cellular health, with a higher ratio indicating less oxidative stress (<u>Owen & Butterfield, 2010</u>). We observed a significantly reduced ratio between control and EtOH exposures in endodermal (20 mM EtOH *P* = 0.007; 70 mM EtOH *P* = 0.006, Student’s *t* test, Fig S4b) and a significantly increased ratio in mesodermal cells (20 mM EtOH *P* = 0.03, Student’s *t* test, Fig. S4b). Even though the GSH/GSSG ratio was not significantly altered between control and EtOH-exposed ectodermal cells, it was significantly lower compared to endodermal (*P* = 3,03×10^−11^, Student’s *t* test) and mesodermal cells (*P* = 1,97×10^−8^, Student’s *t* test).

To predict metabolic pathways affected by EtOH, we performed the metabolite set enrichment analyses (MSEA) with The Small Molecule Pathway Database (SMPDB) library in the MetaboAnalyst 5.0 (<u>Pang et al., 2021</u>) for all identified significantly altered metabolites together and for each germ layer and EtOH concentration separately (Table S16). When all identified metabolites were included in the same analysis, the most significant term was methionine metabolism (Fig. 5e). Methionine metabolism; urea cycle; glutathione metabolism; malate-aspartate shuttle; spermidine and spermine biosynthesis; and aspartate metabolism were significantly altered (*P* < 0.05) in all 70 mM EtOH-exposed germ layers. After the multiple testing correction (FDR < 0.05), alterations in urea cycle; glycine and serine metabolism; and arginine and proline metabolism in the 70 mM EtOH-exposed endodermal and ectodermal cells, as well as methionine metabolism; purine metabolism; and malate-aspartate shuttle in ectodermal cells were observed. However, this pathway analysis underestimates alterations in lipid metabolism and therefore does not give the full picture of alterations.

### Correlations between omics analyses

Finally, to understand the relationship between gene expression, DNAm, and metabolites, relevant correlation analyses were conducted. First, to elucidate the potential effects of DNAm on gene expression, we calculated correlations between 70 mM EtOH-exposed DEGs and the corresponding regulatory region (TSS1500, TSS200, 5’UTR, and 1stExon) probes for each gene separately, and found 111 significant correlations in the endodermal, 18 in the mesodermal, and 78 in the ectodermal cells (*P* < 0.05, Pearson’s correlation coefficient, Table S17a-c). Altogether 36% of the correlating regulatory region probes in the endodermal cells and 42% in the ectodermal cells were hypomethylated and correlated with upregulated genes, including, *LEFTY2* (two probes, *r* = –0.774 and –0.758*, P* = 0.04 and 0.04, respectively), *MIXL1 (r* = –0.9153*, P* = 0.003) and *TAGLN (r* = –0.852*, P* = 0.01) in the endodermal cells as well as *NEUROD4* (three probes, *r* = –0.912, –0.882, and –0.774*, P* = 0.001, 0.003 and 0.02, respectively), *RSPO (r* = –0.782*, P* = 0.02), *NRXN3 (r* = –0.732*, P* = 0.003), and *MAP2 (r* = –0.7844*, P* = 0.02) in the ectodermal cells. Moreover, reduced expression of *LHX2* and *MEG3* correlated with the highest number of probes in the regulatory regions in the ectodermal cells. A total of eight correlated probes of *LHX2* were hypomethylated, and two correlated probes of *MEG3* were hypomethylated and six hypermethylated (Table S17c). Notably, the downregulation of *DLK1*, which is located at the same *DLK1-DIO3* locus as *MEG3*, correlated also with one hypermethylated probe (*r* = –0,838, *P* = 0.009).

Secondly, to clarify the role of metabolism in both DNAm and gene expression, we further tested if the regulatory region probes that significantly correlated with DEGs also correlated with the significantly altered metabolites. The only significant correlations were observed in the ectodermal cells: the hypomethylation of *LHX2* correlated positively with CDP-choline and negatively with LPC 18:1 (18:1/0:0) *(r* = 0.993 and –0.986*, P* = 0.004 and 0.028 after Bonferroni correction, respectively, Pearson’s correlation coefficient), and hypermethylation of *SULF1* correlated negatively with guanosine (*r* = –0.933, *P* < 0.004 after Bonferroni correction, Table S17d).

Thirdly, we calculated correlations between 70 mM EtOH-induced DEGs and significantly altered metabolites (*P* < 0.05), and found four significant correlations in the endodermal, two in the mesodermal, and nine in the ectodermal cells (*P* < 0.05 after Bonferroni correction, Pearson’s correlation coefficient, Table S17e-g). In the ectodermal cells, the most significant correlation was between downregulated *Spermidine/Spermine N1-Acetyltransferase 1* (*SAT1*) and the increased amount of LPC 18:1 (0:0/18:1) (*r* = –0.994, *P* < 0.003). Furthermore, the decreased expression of *DLK1* correlated negatively with LPC 18:1 (0:0/18:1) and LPC 18:1 (18:1/0:0) *(r* = –0.991, *P* = 0.008 and 0.009, respectively), and the decreased expression of *SMOC1* correlated negatively with citric acid (*r* = –0.988, *P* = 0.02).

Finally, we calculated correlations between 70 mM EtOH-induced DMPs and significantly altered metabolites (*P* < 0.05). We observed seven significant correlations in the endodermal, three in the mesodermal, and ten in the ectodermal cells (*P* < 0.05 after Bonferroni correction, Pearson’s correlation coefficient, Table S17h-j). In the endodermal cells, the hypomethylation of *LHX5* correlated negatively with ornithine (*r* = –0.997, *P* = 0.04). In the ectodermal cells, hypomethylation of *LHX2* correlated positively with CDP-choline (*r* = 0.993, *P* = 0.009), hypomethylation of *RAX* correlated negatively with alanine (*r* = –0.991, *P* = 0.02), and hypomethylation of *MEG3* correlated positively with LPC 20:1 (*r* = –0.988, *P* = 0.04).

## DISCUSSION

### EtOH affects important developmental genes and signaling pathways

Here we utilized differentiating germ layer cells to study the effects of EtOH exposure on the transcriptome, methylome, and metabolome in the developmentally crucial gastrulation. We observed several altered genes associated with the major morphogen signaling pathways involved in gastrulation and body patterning, such as WNT, FGF, BMP, the growth and differentiation factor (GDF) as well as transforming growth factor beta (TGF-β) pathways. According to previous research, even small differences in signal duration or intensity during gastrulation can produce different responses, leading to switches in developmental trajectories (<u>Gordon & Blobe 2008</u>). By affecting developmental programming, early PAE could disturb pattern formation and induce a wide spectrum of developmental abnormalities associated with the FASD phenotype. Indeed, it has been shown that a single dose of EtOH in gastrulation produced craniofacial malformations resembling the features of FASD in mouse (<u>Lipinski et al., 2012</u>; <u>Parnell et al., 2009</u>; <u>Sulik et al., 2005</u>).

The largest number of EtOH-induced changes in gene expression, DNAm, and metabolites were observed in the endodermal cells. The majority of the common DEGs (26/30) were more prominently altered in severe EtOH exposure compared to moderate, reflecting a dose response. Among these DEGs was a developmentally essential cardiac hormone *Natriuretic Peptide B* (*NPPB* or *BNP*), which was upregulated in 20 mM EtOH-exposed (effect size 1.2) and even more upregulated in the 70 mM exposed cells (effect size 1.6). A higher amount of this known heart failure biomarker has been observed in the blood samples of heavy alcohol consumers compared to moderate consumers (<u>Zile et</u> <u>al., 2016,</u> <u>Britton et al. 2020</u>), which is in line with our results. Congenital heart diseases including the abnormal development of the heart and great vessels are the most common birth defects worldwide (<u>Zegkos et al., 2019</u>). PAE during the first trimester affects heart development (<u>Carmichael et al. 2003</u>; <u>Tikkanen and Heinonen 1992</u>), and the estimated prevalence of congenital heart defects among individuals with FAS is 40% (<u>Popova et al., 2016</u>). Our findings support the early effects of EtOH on heart development: observed EtOH-induced alterations in endodermal *NPPB, CAT,* and *MAPK1* were detected also recently in adult PAE mice hearts (<u>Atum et al., 2023</u>), and the downregulation of mesodermal *BMP4*, an inducer of early cardiac mesoderm (<u>Tsaytler et al.</u> <u>2023</u>), has also been seen in EtOH-exposed zebrafish embryos (<u>Sarmah et al. 2016</u>). Coordination between the endoderm and mesoderm is crucial for this first developing organ (<u>Van Vliet et al., 2012</u>; <u>Ye et al., 2015</u>), and interestingly, five common DEGs between them, *IGFBP5*, *MIXL1*, *MYL9*, *STC1*, and *TAGLN,* have all functions in the development of heart (<u>Liu et al., 2012</u>; <u>Priya et al., 2020</u>; <u>Zawada et al., 2023</u>).

### EtOH-induced aberrant FGF8 signaling in gastrulation

The most prominent EtOH-induced changes were observed in the ectodermal cells, which is consistent with the previously demonstrated sensitivity of the neurodevelopment to alcohol (<u>Guerri</u> <u>2002;</u> <u>Popova et al., 2023</u>). We observed EtOH-induced downregulation of ectodermal *FGF8* and *SIX3*, as well as upregulation of endodermal *LEFTY1* and *NODAL,* which follow the same altered gene expression patterns of zebrafish homologs when FGF signaling was interrupted. According to zebrafish studies, the lack of FGF signaling induces downregulation of *sixb3,* following by upregulation of *lefty1* and thus disruption of the normal development of central nervous system asymmetry (<u>Gebuijs et al., 2019</u>; <u>Neugebauer & Yost, 2014</u>). Furthermore, we saw downregulation of ectodermal *DUSP6* and upregulation of endodermal *FGF17*, which both have essential roles in the FGF signaling pathway during early embryonic development (<u>Tsukano et al., 2022</u>; <u>Li et al., 2007</u>), and have previously been associated with EtOH exposure in mouse and hESCs (<u>Sun et al., 2023</u>; <u>Khalid et al., 2014</u>). In mice, mutations in *Fgf8* affect particularly the formation of the corpus callosum (<u>Stewart et al., 2016</u>), which enables information transfer in the brain by connecting the left and right cerebral hemispheres (<u>Paul et al., 2007</u>). Notably, based on magnetic resonance imaging studies, it is in particular corpus callosum that is commonly affected in individuals with FASD (<u>Lebel</u> <u>et al., 2011</u>). Our EtOH-induced findings in the FGF regulatory network suggest an early origin for this developmental malformation and elucidate its potential molecular etiology.

Furthermore, *FGF8* and *SIX3* have been found to be mutated in holoprosencephaly (HPE) (<u>Dubourg</u> <u>et al., 2016;</u> <u>McCabe et al., 2011</u>; <u>Wallis et al., 1999</u>), a common developmental defect in midline patterning of the forebrain and/or midface (<u>Dubourg et al., 2007</u>; <u>Marcorelles and Laquerriere 2010</u>). Approximately 1 in 250 embryos is affected by HPE, in which the complex etiology involves both genetic and environmental risk factors (<u>Duborg et al., 2007</u>; <u>Matsunaga et al., 1997</u>). Indeed, based on previous studies, PAE is one of the environmental risk factors for HPE (<u>Abe et al., 2018</u>; <u>Cohen</u> <u>and Shiota, 2002</u>; <u>Croen et al., 2000</u>).

Additionally, we observed ectodermal EtOH-induced alterations in several genes, such as *LHX2*, *SHANK2*, *TRIO*, *ZMYND8*, *ARX*, *NRXN3*, and *SEMA5A*, which have previously been associated also with autism spectrum disorder (ASD) (<u>SFARI Gene Database, 2024</u>). This is in line with the observed overlapping phenotypes between FASD and ASD (<u>Lange et al. 2018</u>) and supports the existence of shared pathogenic mechanisms in these NDDs. Interestingly, haploinsufficiency of *LHX2* has been associated with non-specific NDD phenotype including ASD, variable intellectual disability, speech impairment, microcephaly as well as behavioral, sleep, and brain magnetic resonance imaging abnormalities (<u>Schmid et al. 2023</u>). Since all these features are also linked to FASD phenotype, the role of *LHX2* in the alcohol-induced NDD needs to be elucidated in future studies.

Moreover, significant EtOH-induced alterations in imprinted genes were seen. Due to the essential role of imprinted genes in early development and growth as well as their sensitivity to environmental exposures (<u>Weaver et al., 2009</u>; <u>Kappil et al., 2015</u>), including PAE (<u>Marjonen et al., 2018</u>), ectodermal correlations between hypermethylated probes and downregulation of *DLK1* and *MEG3* in the imprinted *DLK1-DIO3* locus were highly interesting. This locus has previously been associated with both neurodevelopment (<u>Mo et al. 2015</u>, <u>Isles 2022</u>) and PAE in mouse (<u>Laufer et al., 2013</u>). DNAm changes in the imprinted genes were also found in the endodermal cells: the most significantly hypomethylated DMR in *MEST* and hypermethylated DMP in *GNAS*.

### EtOH-induced alterations in metabolites associated with methionine cycle

Consistent with previous results, several significant altered metabolites in this study were associated with methionine metabolism, which provides methyl groups for the synthesis of DNA, polyamines, amino acids, and phospholipids (<u>Ducker et al., 2017</u>; <u>Friso et al., 2017</u>). We observed variable changes between supernatants of the germ layers in different pathways of methionine metabolism, including the methionine, folate and methionine salvage cycles as well as the transsulfuration pathway (Fig. 6). Additionally, nucleosides, nucleotides and their derivatives, several polyamines, amino acids as well as phospholipids were altered, indicating EtOH-induced disturbances in the methionine metabolism. Interestingly, in the severely EtOH-exposed ectodermal cells, both spermidine levels and spermidine acetyltransferase *SAT1* expression was decreased. A polymorphism in *SAT1* has been previously associated with alcohol use disorder (<u>Vaquero-Lorenzo et al., 2014</u>). Spermidine has neuro– and cardioprotective properties (<u>Madeo et al., 2018</u>), and it may mitigate the effects of PAE through its interaction with the N-methyl-D-aspartate (NMDA) glutamate receptors (<u>Honse et al., 2003</u>). EtOH-induced decreased binding activity of NMDA receptors can disrupt neurodevelopment associated with PAE in mouse (<u>Honse et al., 2003</u>; <u>Savage et al., 1991</u>). We also observed significantly decreased amounts of neurotransmitters glutamic acid and acetylaspartylglutamic acid in ectodermal cells, which is in line with EtOH-induced disruptions of NMDA receptors.

**Fig. 6.**
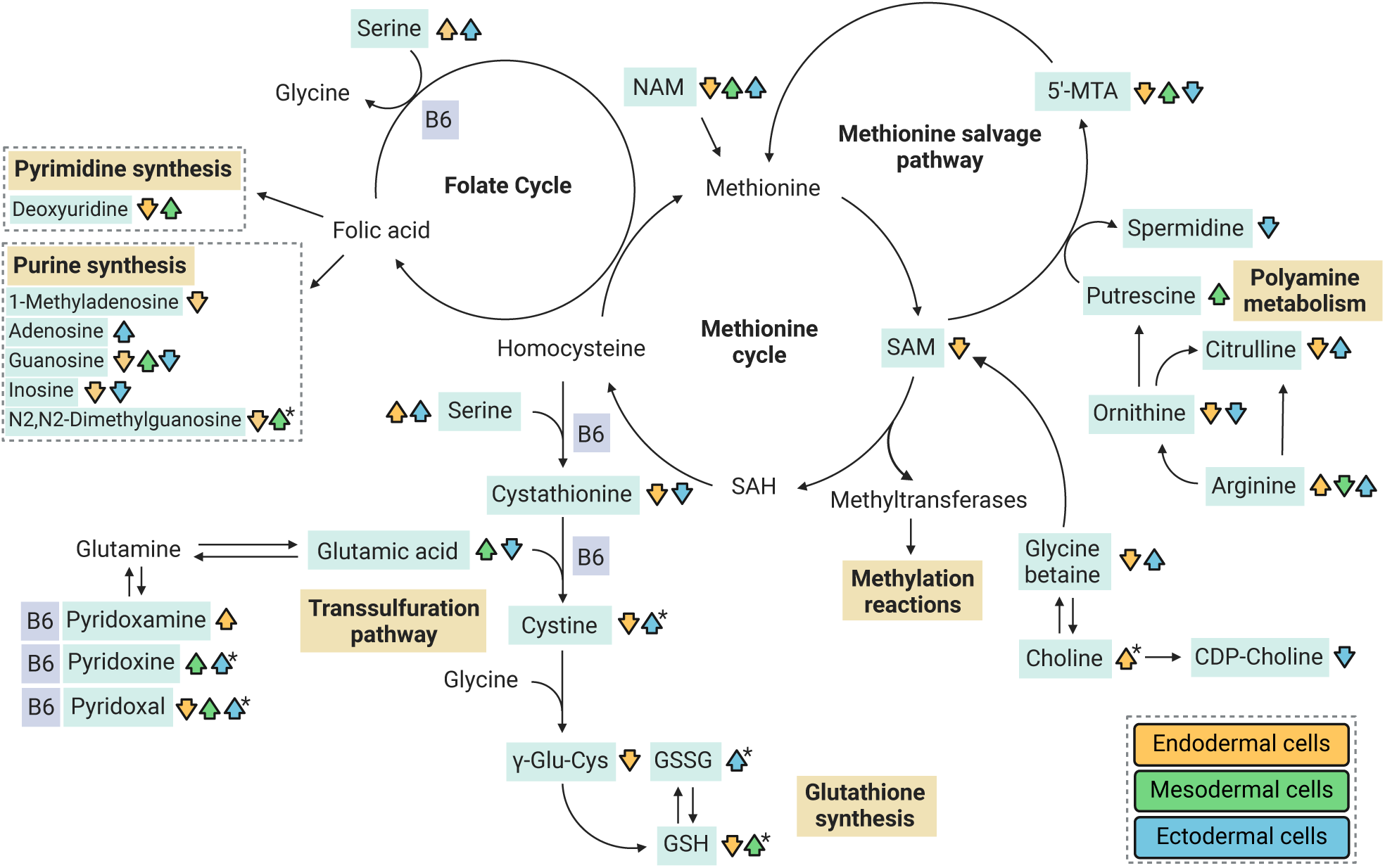
EtOH-induced alterations in the metabolites of the methionine metabolism. Arrows mark significant EtOH-induced alterations in both exposures or only in 70 mM EtOH exposure. Arrow with one star mark significant EtOH-induced alterations only in 20 mM EtOH exposure. 5’-MTA, 5’-methylthioadenosine; B6, Vitamin B6; GSH, reduced glutathione; GSSG, oxidised glutathione; NAM, N-acetylmethionine; SAM, S-adenosylmethionine; γ-Glu-Cys, γ-glutamyl-cysteine. Created in Biorender.

Previous studies have shown that methyl donors, betaine, choline, folic acid, and B6 vitamins, can play a protective role against PAE-induced epigenetic changes and adverse effects on the phenotype (<u>Bekdash et al., 2013</u>; <u>Karunamuni et al., 2017</u>; <u>Jacobson et al., 2018</u>). Although the directions of EtOH-induced alterations in methyl donors varied between the germ layers, and the culture medium contains an excess amounts of methyl donors, there were still alterations in choline, CDP-choline, glycine betaine and different forms of B6 vitamin. We also observed reduced levels of endodermal and ectodermal NAD+ and downregulation of endodermal *NMRK1*, which encodes nicotinamide riboside kinase that acts as NAD+’s precursor. NAD+ has several important roles in methionine metabolism by serving as a cofactor for key enzymes, regulating methylation reactions, influencing polyamine synthesis, and homocysteine recycling (<u>Ducker et al., 2017;</u> <u>Covarrubias et al., 2021;</u> <u>Wang et al., 2022</u>). Moreover, NAD+ acts as a coenzyme in EtOH metabolism (<u>Zakhari et al., 2006</u>).

### Effects of EtOH exposure on gene regulation

The effects of EtOH on methionine cycle and methylation potential is supported by the observed genome-wide hypomethylation in relation to islands in the mesodermal cells as well as in the regulatory regions in the ectodermal cells. Also, in the endodermal and ectodermal cells, a notable number of upregulated DEGs were correlated with hypomethylated probes in the regulatory regions, suggesting effects of EtOH on gene regulation. In contrast to our recent hESCs work (<u>Auvinen et al.</u> <u>2022</u>), we did not detect decreased expression of DNA methyltransferases *DNMT3A* or *DNMT3B* in the current study. Interestingly, despite the lower SAM/SAH ratio in the most altered endodermal cells, no EtOH-induced changes were observed in methionine or SAH. However, i*n vivo* conditions are very different, and disrupted methionine cycle combined with poor maternal nutrition (e.g. folate) can cause much larger changes in cellular methylation potential than we have seen in this *in vitro* work.

In addition to DNAm, PAE has been found to change the regulation of fetal genes through histone acetylation in mouse (<u>Mews et al., 2019</u>). This is in line with previous studies, in which the EtOH-induced alterations in gene expression involved in mouse cardiac development was caused by increased histone acetylation, not by altered DNAm (<u>Choudhury & Shukla, 2008</u>; <u>Kim & Shukla,</u> <u>2006</u>), also *in vitro* (<u>Zhong et al. 2010</u>). Furthermore, in our work, one metabolite and several genes associated with the molecular mechanisms of acetylation, such as N-acetyllysine, *LNCPRESS1, CDK2AP1, ACAT2,* and *SALL1,* were altered. This clearly indicates that there are epigenetic mechanisms other than DNAm and these, like histone acetylation, other histone modifications, and small RNAs should be considered in future studies.

### Limitations of the study and future prospects

This *in vitro* model enables examination of the effects of EtOH exposure on human cells under controlled conditions and without confounding factors in a challenging developmental period in human research. We used the H1 line, which consists of well-characterized and extensively studied reference hESCs. However, by using only one cell line, we cannot determine the effects of genetic background on the observed alcohol-induced changes. Furthermore, in this relatively simple model, it is not possible to determine the causality or stability of the changes or obtain information about the effects of EtOH on the interactions of the germ layers and the further development. Also, by using extracellular supernatants, we were only able to study the extracellular metabolites. However, many genes and metabolites which were altered by EtOH in the current study have also been associated with PAE or EtOH exposure in different organisms in several *in vivo* and *in vitro* studies previously, supporting our findings. The results that we present are associations and first steps towards understanding the effects of early PAE and the etiology of the variable FASD phenotype.

In addition to elucidating the etiology of alcohol-induced developmental disorders, our aim was to examine the sensitivity of gastrulation to alcohol exposure. Due to a large proportion (44–65%) of unplanned pregnancies (<u>Bearak et al., 2018</u>), there is a considerable risk of PAE prior to pregnancy recognition (<u>Voutilainen et al., 2022</u>). In the current study, both moderate (20 mM) and severe (70 mM) EtOH exposures induced changes in gene function, although the moderate exposure induced statistically significant alterations only in the endodermal cells. However, based on these results, we do not know how much EtOH is required for biologically significant changes during gastrulation. Furthermore, although the peak EtOH concentration of culture medium in the moderate exposure corresponded approximately 0.9‰, and the severe EtOH exposure about 3.2‰, the effects of *in vitro* and *in vivo* exposures cannot be directly compared. Considering the most prominent EtOH-induced changes in the ectodermal cells in the current work as well as the sensitivity of neurodevelopment to EtOH observed in previous studies, the possibility of NDD caused by early PAE needs to be carefully investigated. Future studies using advanced *in vitro* methods such as embryo models and human forebrain organoids as well as appropriate human cohorts will clarify the effects of early PAE on human brain development and consequently on the phenotype.

## METHODS

### Maintenance and differentiation of hESCs and EtOH exposure

*hESC culture.* hESC cell line H1 (WA01) was obtained from Biomedicum Stem Cell Center (BSCC, Helsinki, Finland) through a license agreement with WiCell, Inc. hESCs were cultured in E8 Medium (Gibco) on Matrigel (Corning) coated plates at 37 °C and 5% CO2. Culture media was routinely replaced every day, and cells were passaged using 0.5 mM EDTA.

*Germ layer cell differentiation.* hESCs were differentiated into the three germ layers by using the STEMdiff Trilineage Differentiation kit (Stemcell Technologies). Single-cell suspensions were prepared according to the manufacturer’s instructions with 10 μM Y-27632 for the first 24 h after seeding. Cells were seeded in 6-well plates at the following densities: 250,000 cells/well for the mesodermal, 1 million cells/well for the endodermal, and 1.5 million cells/well for the ectodermal cells. Differentiation was performed according to the manufacturer’s instructions, and the medium was changed daily. Endodermal and mesodermal cells were collected after five days and ectodermal cells after seven days.

*EtOH exposure.* The medium was supplemented with EtOH (≥ 99.5 p-%) at a final concentration of 20 mM or 70 mM. Exposure was started at the same time as the differentiation (day 1) and exposure times were four days (96 h) for endodermal and mesodermal cells and six days (144h) for ectodermal cells. The experiment was performed simultaneously with four or five replicates per germ layer.

### DNA and RNA extractions

DNA and RNA were extracted simultaneously using AllPrep DNA/RNA/miRNA Universal Kit (Qiagen) according to the manufacturer’s instructions. RNA quality was assessed using an Agilent 2100 Bioanalyzer (Agilent Technologies, Inc.), which was provided by the Biomedicum Functional Genomics Unit (FuGU) at the Helsinki Institute of Life Science and Biocenter Finland at the University of Helsinki.

### 3’mRNA sequencing (mRNA-seq) analysis

*Differential expression analysis.* The extracted RNA was prepared by diluting samples to 10 ng/μl and the library preparation for bulk 3’-sequencing of poly(A)-RNA was performed as previously described (<u>Macosko et al. 2015</u>) provided by FuGU. The libraries were sequenced on NextSeq 500 (Illumina) with three separate batches. Drop-seq pipeline was used to construct the mRNA-seq count table for RNA samples. 36,087 transcripts were identified for downstream analysis. The DESeq2 R package (<u>Love et al. 2014</u>) was used to detect the DEGs (FDR-corrected *P*-value < 0.05) separately for each germ layer with the model adjusting for batch. Benjamini-Hochberg procedure was used to control the FDR. Principal component analysis identified one batch with three outlier samples, which were removed from the analysis. Volcano plots and heat maps were plotted using ggplot2 and pheatmap R packages.

*Pathway analysis. enrichgo* function in R package clusterProfiler version 4.8.2. (<u>Wu et al. 2021</u>) was used to perform gene-set enrichment analysis for DEGs. The GO and KEGG knowledgebases were used as the source for identifying significantly enriched BP, CC, MF, and KEGG terms (FDR-corrected *q*-value < 0.05). Benjamini-Hochberg procedure was used to control the FDR. Due to the low number of DEGs, pathway analysis was not performed for mesodermal cells.

### Validation of differentiation

To verify the identity of the established germ layers, we evaluated expression levels of key germ layer marker genes by 3’mRNA sequencing. Unexposed hESCs and germ layer cells exhibited characteristic expression levels for each cell type (Fig. S5), which confirmed a successful trilineage differentiation. Further characterization with immunofluorescence analysis illustrated key germ layer marker expression in each germ layer (Fig. S6). The undifferentiated hESCs and differentiating stem cells were seeded onto 24-well culture plates with 30,000 cells per well before the immunostainings. Cells were fixed in 4% paraformaldehyde (PFA, Fisher Chemical) in PBS for 10 min and washed with PBS. Immunostaining was performed with antibodies for SOX17 (R&D systems AF1924, Goat) in endodermal, NCAM (EMD Millipore AB5032, Rabbit) in mesodermal, and PAX6 (Thermo Fisher Scientific 42-6600, Rabbit) in ectodermal cells. Cells were imaged with EVOS FL Cell Imaging System (Thermo Fisher Scientific).

### DNAm microarrays

*Genome-wide DNAm analysis.* Genomic DNA (1,000 ng) was sodium bisulfite-converted using the Zymo EZ DNAm™ kit (Zymo Research) and genome-wide DNAm was assessed with Infinium Methylation EPIC BeadChip v1.0 (Illumina, San Diego, CA, USA) following standard protocol at the Institute for Molecular Medicine Finland. The raw DNAm dataset was pre-processed, quality-controlled, and filtered with the ChAMP R package (<u>Tian et al. 2017</u>) using the minfi method (<u>Fortin</u> <u>et al., 2017</u>). Data was filtered using detection *P*-values and bead count values with default thresholds. Probes were normalized with noob (<u>Triche et al., 2013</u>) in minfi R package (<u>Fortin et al., 2017</u>), and BMIQ in wateRmelon R package (<u>Pidsley et al., 2013</u>). No batch effect correction was needed for the data. After adjustment, probes located in sex chromosomes and probes binding to polymorphic and off-target sites were filtered. After filtering steps, 756 351 probes were retained for further downstream analysis. Annotation information was merged to corresponding probes from IlluminaHumanMethylationEPICanno.ilm10b4.hg19 R package (<u>Hansen, 2017</u>), which is based on the file “MethylationEPIC_v-1-0_B4.csv” from Illumina. The following abbreviations were used: TSS1500, TSS200, 5’UTR, 3’UTR, N_shelf: north shelf, N_shore, S_shore, S_shelf: south shelf. Probes were annotated to genes based on UCSC and GENCODE Comprehensive V12 (GencodeComp) databases. The names of genes were used primarily according to UCSC, and the location of genes were primarily determined according to GencodeComp.

*GWAM analysis.* β-values of all normalized probes in the array were used to calculate GWAM levels sample-wise (<u>Li et al., 2018</u>). Differences between EtOH-exposed and control samples were calculated by selecting the appropriate statistical test from two-tailed Student’s t-test, Welch Two Sample t-test, or Wilcoxon rank sum exact test based on normality and variances. Normality of the data was assessed with Shapiro-Wilk normality test and the variances were compared with F-test. Gene locations were used according to UCSC. If the gene location information was missing, probe was marked as “unknown.” In the case of multiple location entries, group “others” was used.

*RE DNAm analysis.* Processed DNAm data (M values) was used to predict DNAm in Alu, ERV, and LINE1 elements sample-wise using a Random Forest-based algorithm implemented by REMP R package (<u>Zheng et al., 2017</u>). Less reliable predicted results were trimmed according to the default quality score threshold 1.7 and missing rate 0.2 (20%). Differences between EtOH-exposed and control samples were calculated by selecting the appropriate statistical test from two-tailed Student’s t-test, Welch Two Sample t-test or Wilcoxon rank sum exact test based on normality and variances. Normality of the data was assessed with Shapiro-Wilk normality test and the variances were compared with F-test.

*DMP analysis.* DMP analysis was performed by Limma R package (<u>Ritchie et al., 2015</u>) by using M-values. β-values were used for visualization and interpretation of the results. DMPs were considered significant when the FDR-corrected *P*-value was < 0.05. Benjamini-Hochberg procedure was used to control FDR. Volcano plots were plotted using ggplot2 R packages.

*DMR analysis.* DMRcate R package was used for analyzing DMRs (<u>Kolde et al. 2016</u>). In short, analysis in DMRcate begins by standard linear modeling, which produces t values for each CpG site. DMRcate then applies kernel smoothing, which factors in the neighboring sites, giving weighted average values for each CpG site. Model for the smoothed data is then created and the CpG sites with adjusted *P*-values below the threshold are agglomerated into regions. DMRcate was adjusted to determine probes (≥ 3) in a region with a maximal allowed genomic distance of 1000 bp, and scaling factor C was set at 2, as per the authors’ recommendation (<u>Kolde et al. 2016</u>). Default *P*-value cutoffs, including FDR < 0.05 for individually significant CpGs, were used in the analysis.

*Pathway analyses.* Enrichment analysis was performed for significant DMPs (*P* < 0.05) by *gometh* function in missMethyl R package (<u>Phipson et al. 2015</u>), which considers the different number of probes per gene present on the EPIC array and CpGs that are annotated to multiple genes. missMethyl was set to use the GO database as the source for identifying significantly enriched BP, CC, and MF terms and KEGG database from genes that contained at least one significant DMP. Terms with nominal *P* < 0.05 were reported as only the terms of endodermal cells were significant with FDR correction. Due to the low number of DMPs in mesodermal cells and DMRs in all germ layer cells, they were not subjected to pathway analyses.

### Metabolomic analysis

*Sample processing.* Metabolomics analysis was conducted on the supernatants of the same samples as in the DNAm and gene expression analysis. For the metabolite extraction, the samples were randomized, and 400 µL of cold methanol (100%) was added to 100 µL of sample. The samples were kept in ice between the steps. The pooled quality control (QC) sample was prepared by combining 10 µL from each sample. The supernatants were filtered (Acrodisc 4 mm with 0.45 μm membrane) and inserted into HPLC vials for analysis.

*LC–MS metabolite profiling analysis.* The samples were analyzed by liquid chromatography–mass spectrometry (LC-MS), consisting of an ultra-high performance liquid chromatography (TUPLC) combined with Thermo Q Exactive^TM^ Hybrid Quadrupole-Orbitrap mass spectrometer (Thermo Scientific). The technical specifications of the reverse phase (RP) column were: Zorbax Eclipse XDB-C18, particle size 1.8 µm, 2.1 × 100 mm (Agilent Technologies). For the hydrophilic interaction chromatography (HILIC) column, the technical specifications were: Acquity UPLC BEH Amide 1.7 µm, 2.1 × 100 mm (Waters Corporation). The chromatography parameters were as follows: for RP chromatography, the column oven temperature was set to 50°C, and the flow rate to 0.4 mL/min gradient elution with water (eluent A) and methanol (eluent B) both containing 0.1% (v/v) of formic acid. Gradient profile for RP separations was 0 to 10 min: 2% B → 100% B; 10 to 14.5 min: 100%

B; 14.5 to 14.51 min: 100% B → 2% B; 14.51 to 16.5 min: 2% B. For HILIC, the column oven temperature was set to 45°C, flow rate 0.6 ml/min, gradient elution with 50% v/v ACN in water (eluent A) and 90% v/v ACN in water (eluent B), both containing 20 mM ammonium formate (pH 3). The gradient profile for HILIC separations was 0 to 2.5 min: 100% B, 2.5 to 10 min: 100% B → 0% B; 10 to 10.01 min: 0% B → 100% B; 10.01 to 12.5 min: 100% B. Both negative and positive electrospray ionization (ESI) were used for both analytical modes. ESI settings were ray voltage 3.5 kV for positive and 3.0 kV for negative, max spray current 100, flow rates 40 for sheath gas, 10 for auxiliary gas, and 2 for spare gas (as arbitrary units for ion source), S-lens RF level 50 V, capillary and probe heater temperature 300 °C. A full scan range from m/z 60 to 700 for HILIC modes and 120 to 1200 for RP modes with the resolution of 70 000 (m/Δm, full width at half maximum at 200 u) and an automated injection time and gain control targeted at 1 000 000 ions. For tandem mass spectrometry (MS/MS), three peaks with apex trigger 0.2 to 3 s were selected for MS/MS fragmentation with 15 s dynamic exclusion. Normalized collision energy at 20, 30, and 40%, was used in MS/MS with a mass resolution of 17 500 (m/Δm, full width at half maximum at 200 u), an automated gain targeted at 50 000, and isolation window 1.5 m/z.

*Data analysis.* Peak detection and alignment were performed in MS-DIAL ver 4.90 (<u>Tsugawa et al.</u> <u>2020</u>). For the peak collection full scale of m/z values for each mode were considered and retention times after 0.5 min were considered. The amplitude of the minimum peak height was set at 300 000 in the negative modes or 500 000 in the positive modes. The peaks were detected using the linear weighted moving average algorithm. For the alignment of the peaks across samples, the retention time tolerance was 0.1 min and the m/z tolerance was 0.01 Da. Drift correction and removal of low-quality signals were done with Notame R-package as described in Klåvus et al. (<u>2020</u>). For feature-wise analysis, we used Welch’s t-test and Cohen’s d-effect sizes.

*Compound identification.* The chromatographic and mass spectrometric characteristics (retention time, exact mass, and MS/MS spectra) of the significantly differential molecular features were compared with entries in an in-house standard library and publicly available databases, such as METLIN and The Human Metabolome Database (HMDB), as well as with published literature. The annotation of each metabolite and the level of identification was given based on the recommendations published by the Chemical Analysis Working Group (CAWG) Metabolomics Standards Initiative (MSI) (<u>Sumner et al., 2007</u>).

*Pathway analyses.* MetaboAnalyst 5.0 was used for the metabolite set enrichment analyses (MSEA) with The Small Molecule Pathway Database (SMPDB) library of the annotated metabolites (<u>Pang et</u> <u>al., 2021</u>). Enrichment analysis was performed for the metabolites, which were significantly altered by EtOH (*P* < 0.05) with an available HMDB ID as follows: 40 metabolites in 20 mM EtOH-exposed endodermal cells, 33 metabolites in 70 mM endodermal cells, 15 metabolites in 20 mM mesodermal cells, 27 metabolites in 70 mM mesodermal cells, 21 metabolites in 20 mM ectodermal cells, 55 metabolites in 70 mM ectodermal cells, and 79 metabolites in all germ layer cells together.

### Correlation analysis

Correlations between DEGs and CpGs in the corresponding regulatory region for each gene, DEGs and differentially altered annotated metabolites, as well as DMPs and differentially altered annotated metabolites were calculated with Pearson correlation. β-values of CpGs, variance stabilizing transformed (vst) expression counts and signal intensities of molecular features were used. In the analysis, CpGs annotated only for TSS1500, TSS200, 5’UTR and 1stExon regions, based on GencodeComp annotation, were considered as regulatory region CpGs. Analysis for correlation between DEGs and the corresponding regulatory region CpGs was calculated for each gene separately, and thus unadjusted *P*-values were used to determine the significance of results. Bonferroni correction was used for multiple testing correction in all other analyses due to a high number of comparisons.

## DATA AVAILABILITY AND ACCESS

All relevant data can be found within the article and its supplementary information.

## SUPPLEMENTARY MATERIAL

Supplementary Figures 1-6

Supplementary Tables 1-17

