## Supplementary Figures for "Effects of alcohol on the transcriptome, methylome, and metabolome of *in vitro* gastrulating human embryonic cells"

**Figure S1:** GWAM comparison between controls and 70 mM EtOH-exposed germ layer cells.

**Figure S2:** Comparison of RE DNAm between controls and 70 mM EtOH-exposed germ layer cells.

**Figure S3:** Heat map of the normalized abundances of all annotated metabolites in the germ layers.

**Figure S4:** Ratios of SAM/SAH and GSH/GSSG in the germ layers.

**Figure S5:** Validation of hESC differentiation into the germ layer cells by gene expressi.

**Figure S6:** Validation of hESC differentiation into the germ layer cells by immunofluorescence staining.

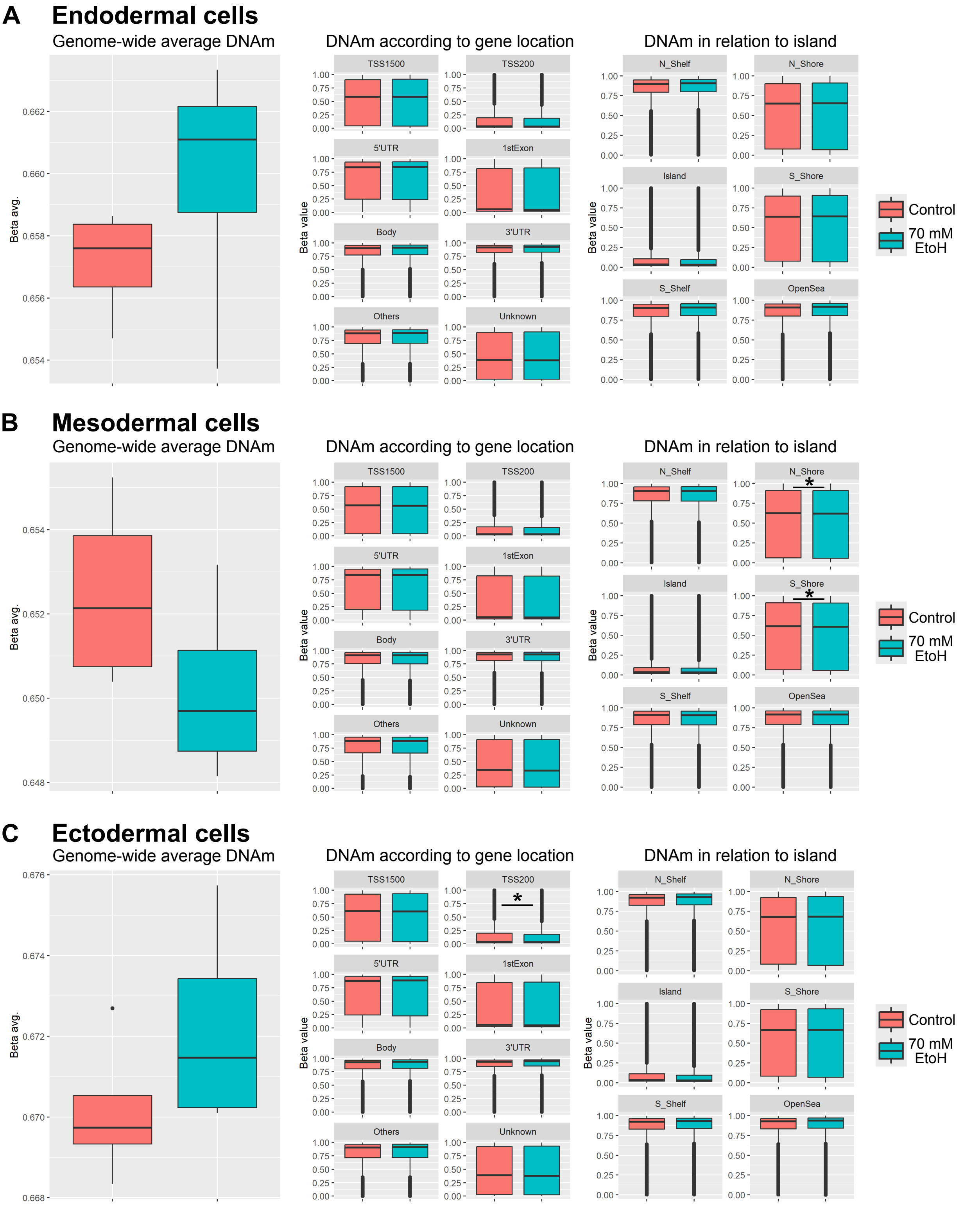


**Supplementary figure 1: GWAM comparison between controls and 70 mM EtOH-exposed germ layer cells.** Comparison of DNAm in all probes, in relation to gene, and in relation CpG island in **a** endodermal cells, **b** mesodermal cells, and **c** ectodermal cells. **P* < 0.05, Student's t test or Welch two sample t test. TSS1500: 1500 bp upstream of transcription start site, TSS200: 200 bp upstream of TSS, UTR: untranslated region, N_shelf: north shelf, N_shore: north shore, S_shore: south shore, S_shelf: south shelf.

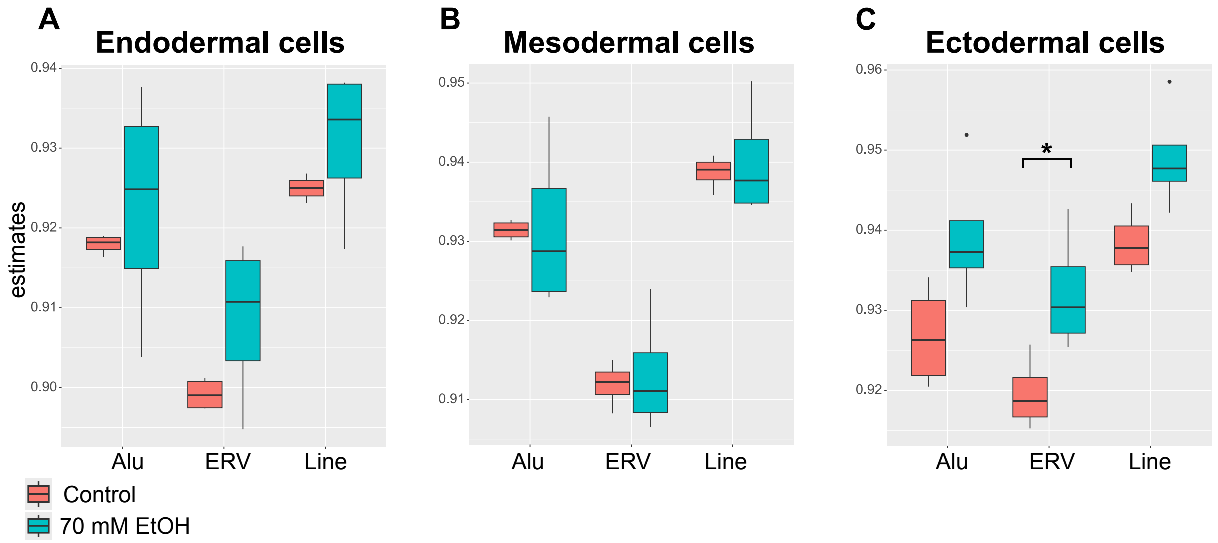


**Supplementary figure 2: Comparison of RE DNAm between controls and 70 mM EtOH-exposed germ layer cells.** Comparison of DNAm in Alu, ERV, LINE1 repetitive region in a endodermal cells, b mesodermal cells, and c ectodermal cells. * P < 0.05, Student's t test.

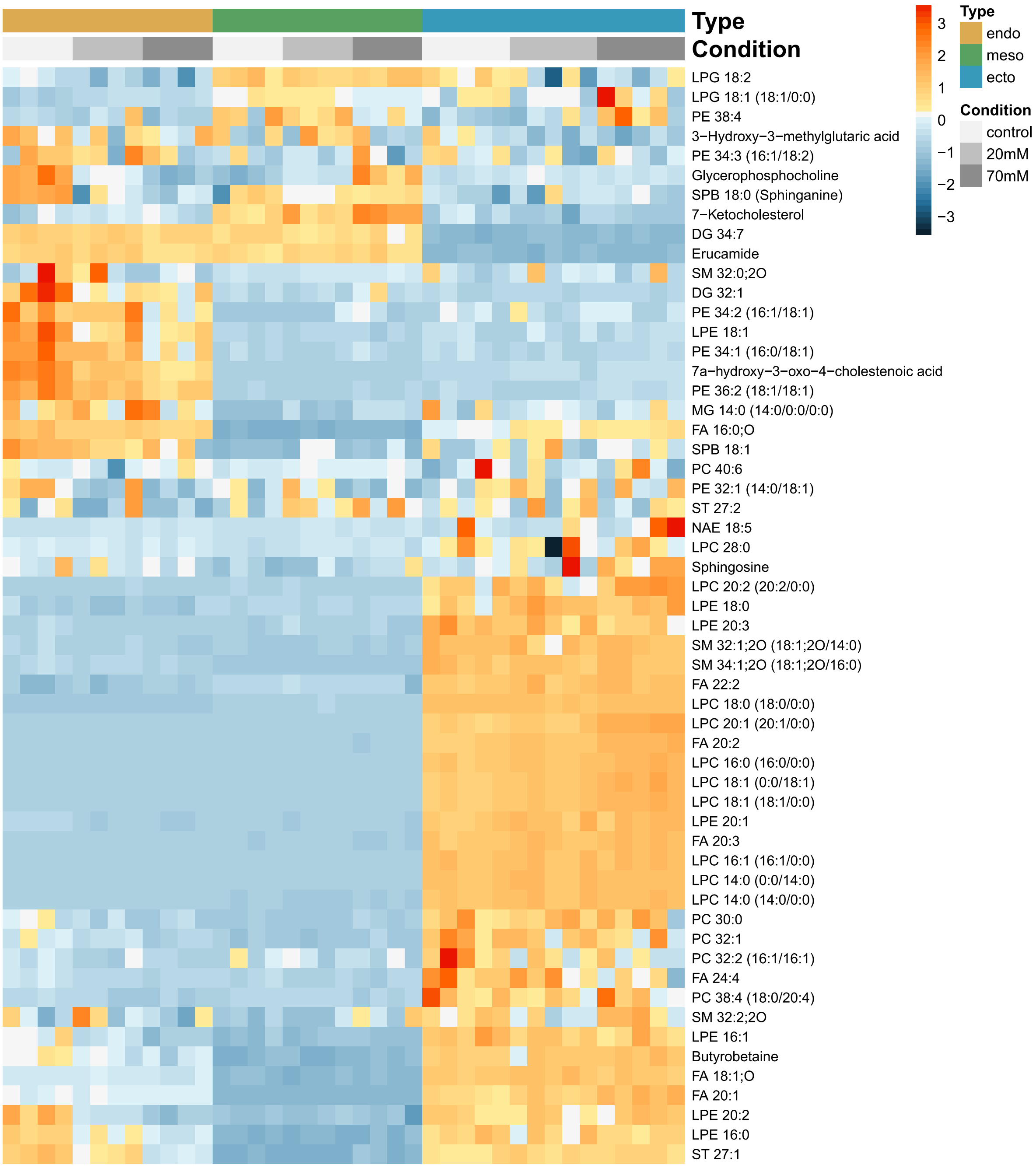
Type
Condition
LPG 18:2
LPG 18:1 (18:1/0:0)
PE 38:4
3−Hydroxy−3−methylglutaric acid
PE 34:3 (16:1/18:2)
Glycerophosphocholine
SPB 18:0 (Sphinganine)
7−Ketocholesterol
DG 34:7
Erucamide
SM 32:0;2O
DG 32:1
PE 34:2 (16:1/18:1)
LPE 18:1
PE 34:1 (16:0/18:1)
7a−hydroxy−3−oxo−4−cholestenoic acid
PE 36:2 (18:1/18:1)
MG 14:0 (14:0/0:0/0:0)
FA 16:0;O
SPB 18:1
PC 40:6
PE 32:1 (14:0/18:1)
ST 27:2
NAE 18:5
LPC 28:0
Sphingosine
LPC 20:2 (20:2/0:0)
LPE 18:0
LPE 20:3
SM 32:1;2O (18:1;2O/14:0)
SM 34:1;2O (18:1;2O/16:0)
FA 22:2
LPC 18:0 (18:0/0:0)
LPC 20:1 (20:1/0:0)
FA 20:2
LPC 16:0 (16:0/0:0)
LPC 18:1 (0:0/18:1)
LPC 18:1 (18:1/0:0)
LPE 20:1
FA 20:3
LPC 16:1 (16:1/0:0)
LPC 14:0 (0:0/14:0)
LPC 14:0 (14:0/0:0)
PC 30:0
PC 32:1
PC 32:2 (16:1/16:1)
FA 24:4
PC 38:4 (18:0/20:4)
SM 32:2;2O
LPE 16:1
Butyrobetaine
FA 18:1;O
FA 20:1
LPE 20:2
LPE 16:0
ST 27:1
3
2
1
0
−1
−2
−3
Type
endo
meso
ecto
Condition
control
20mM
70mM

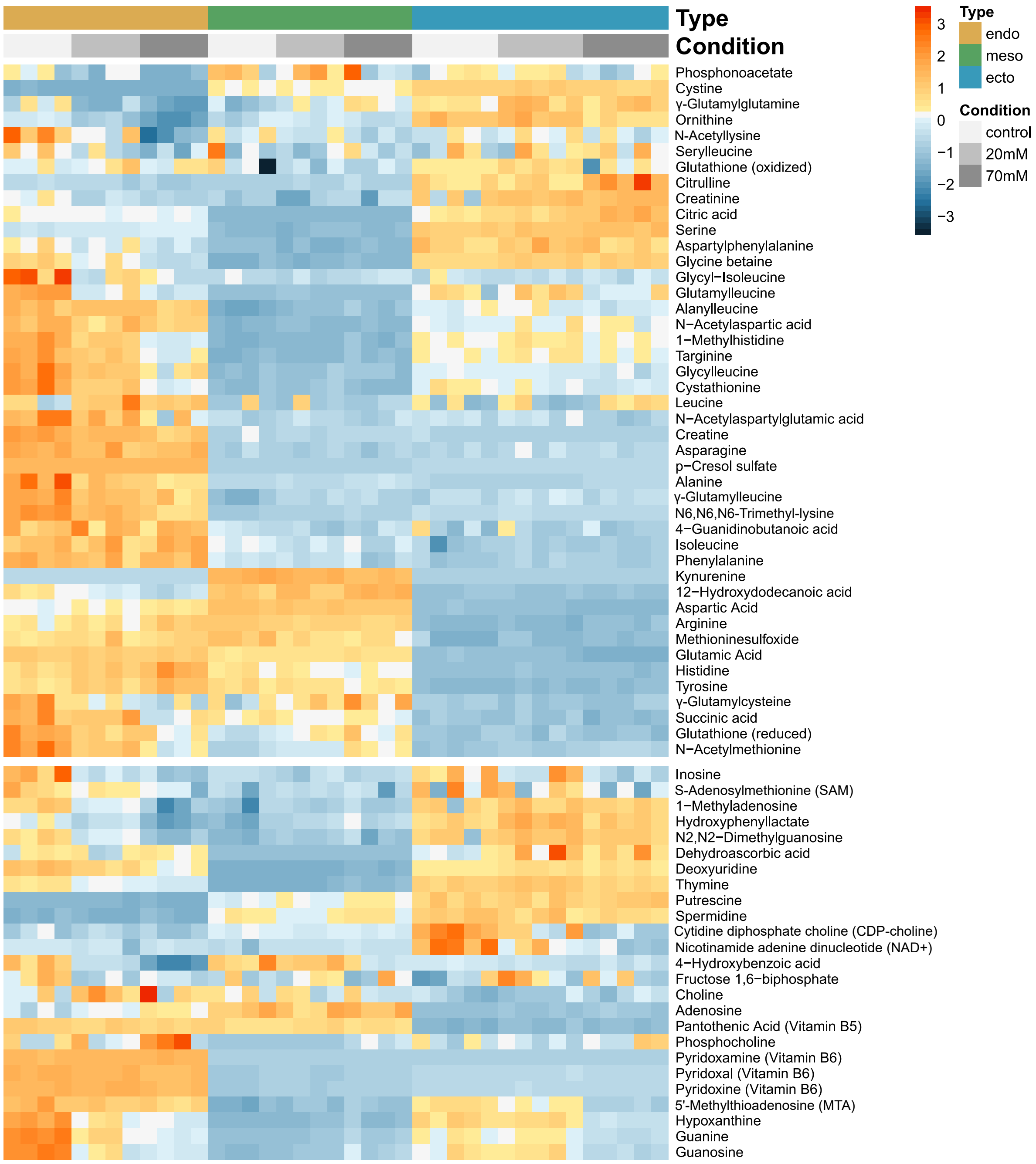


**Supplementary figure 3: Heat map of the normalized abundances of all annotated metabolites in the germ layers.** Hierarchical clustering was applied to arrange the metabolites based on their similarity of the abundance between the samples.

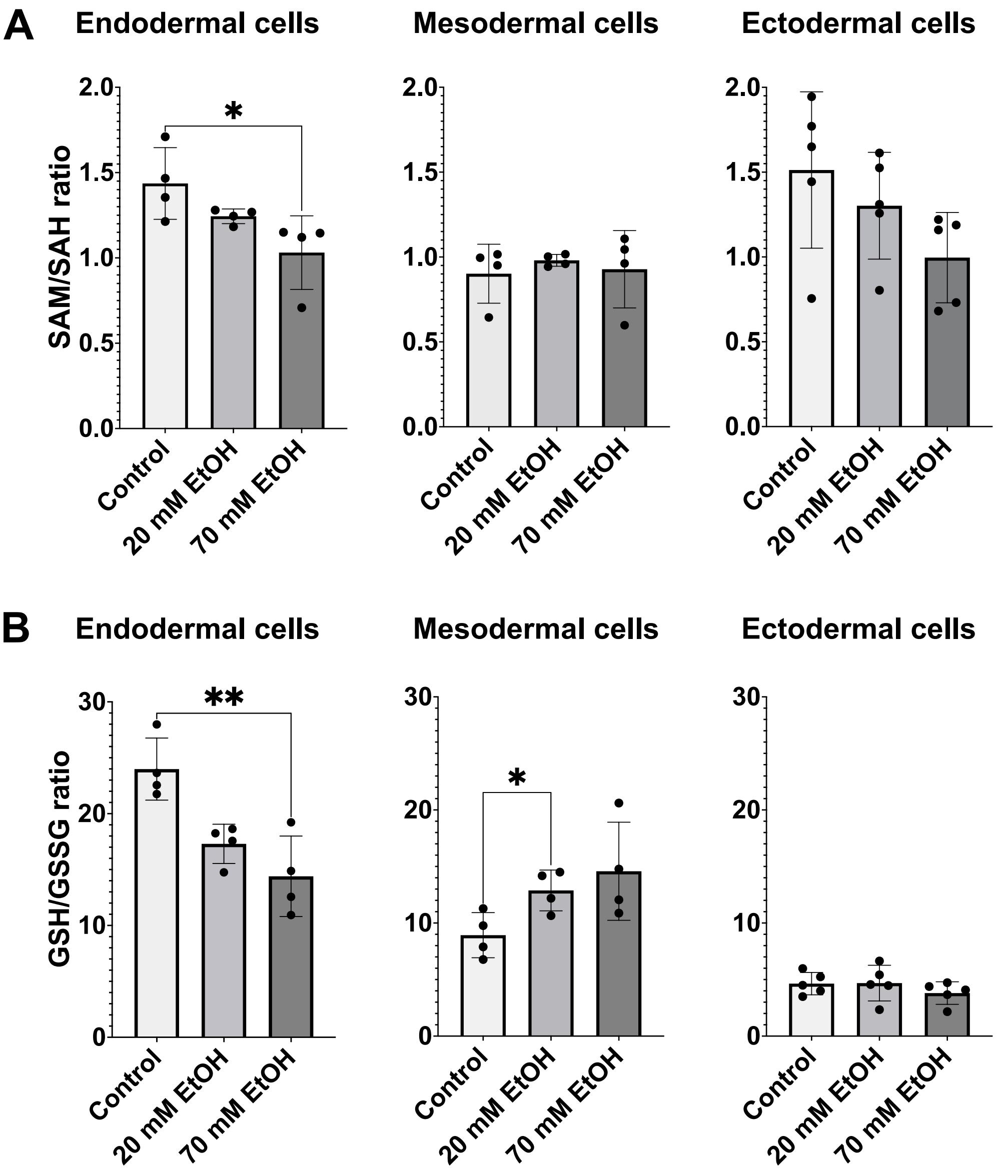


**Supplementary figure 4: Ratios of SAM/SAH and GSH/GSSG in the germ layers.** Ratios of S-adenosylmethione (SAM) and S-adenosylhomocysteine (SAH) as well as reduced glutathione (GSH) and oxidised glutathione (GSSG). Ratios presented as mean ±SD. Endodermal and mesodermal cells $n = 4$, ectodermal cells $n = 5$. *$P$-value < 0.05, **$P$-value < 0.01.

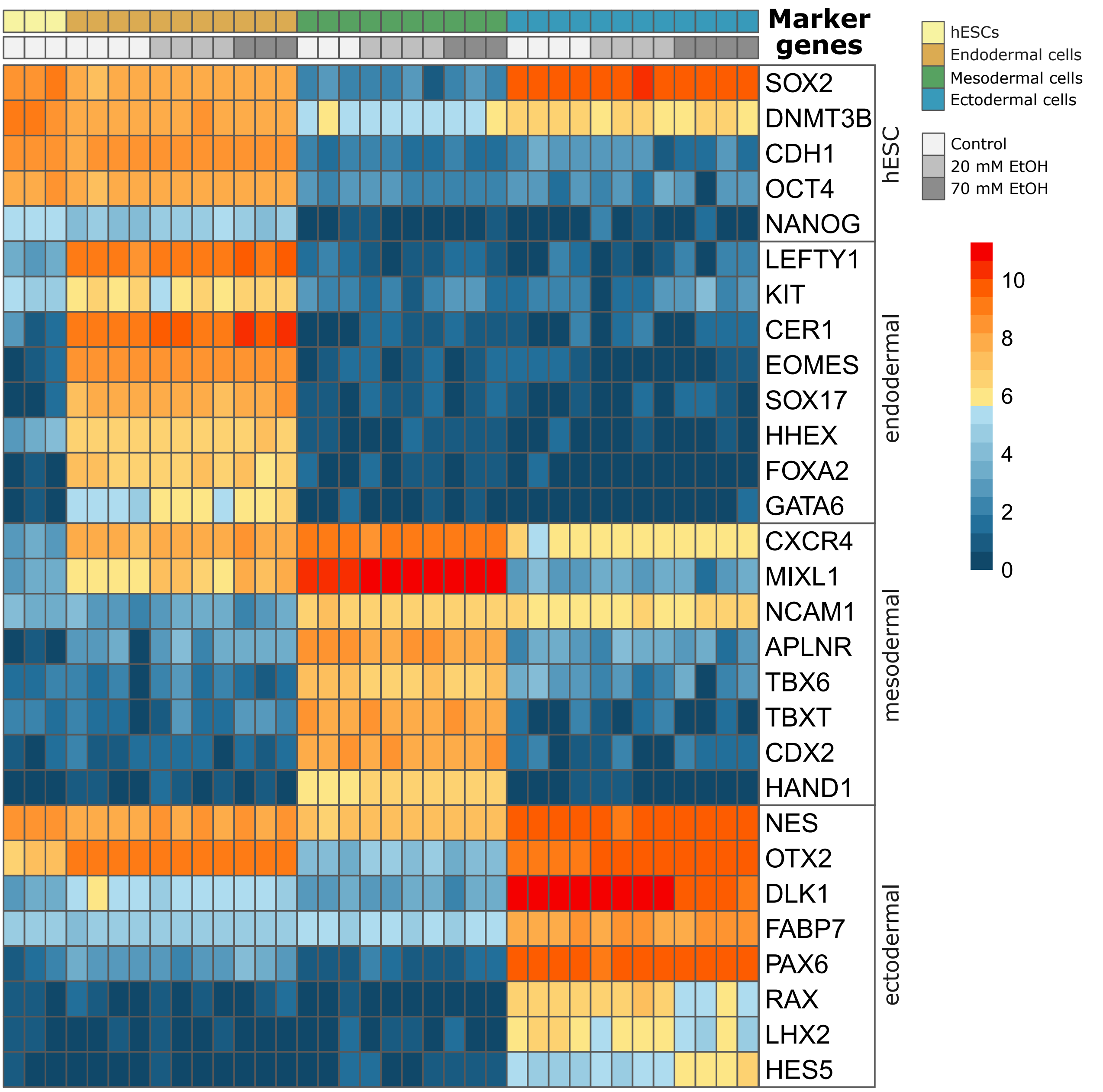


**Supplementary figure 5: Validation of hESC differentiation into the germ layer cells by gene expression.** Gene expression heatmap of pluripotency and differentiation marker genes in hESCs as well as endodermal, mesodermal and ectodermal cells.

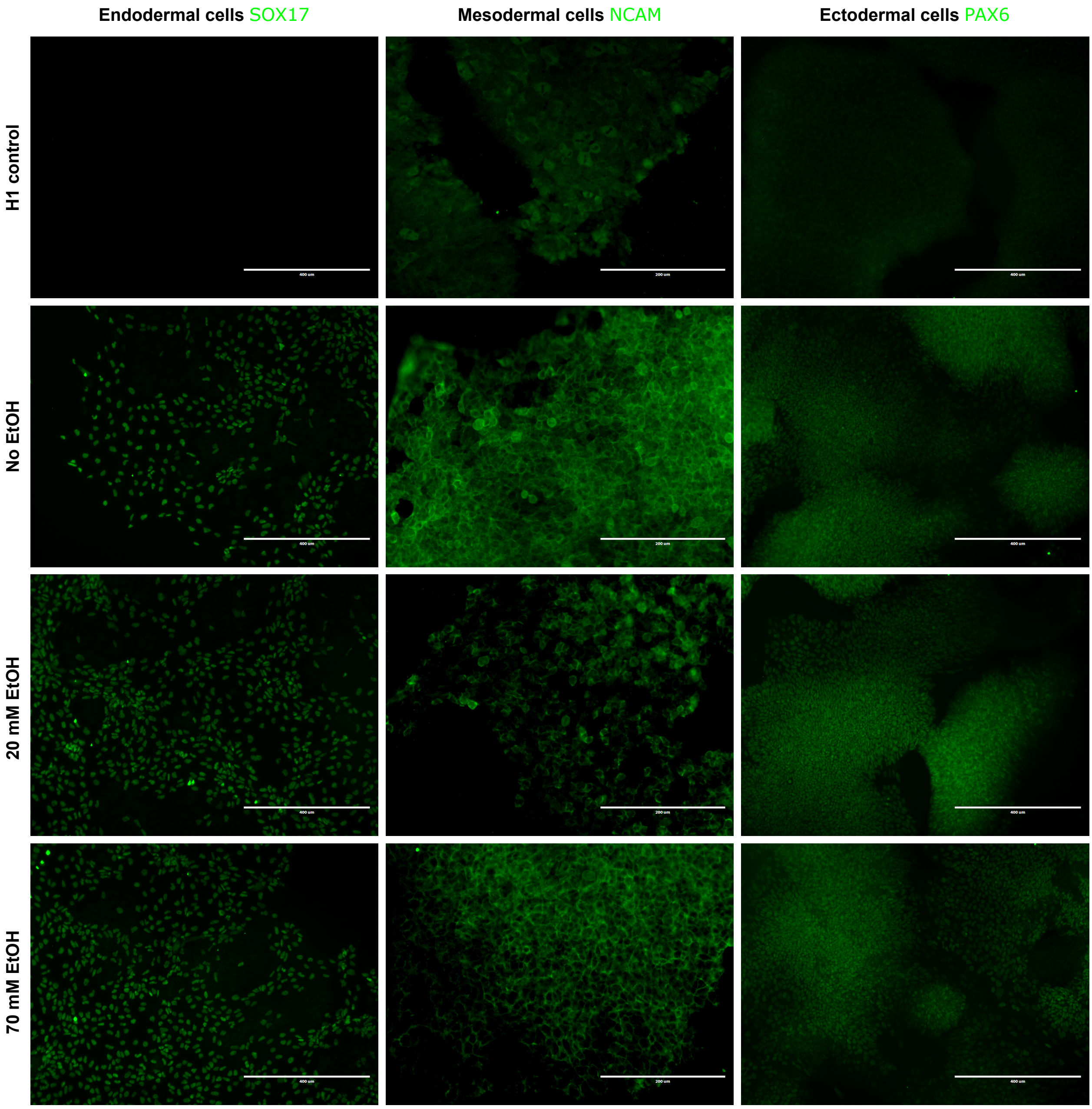


**Supplementary figure 6: Validation of hESC differentiation into the germ layer cells by immunofluorescence staining.** Immunofluorescence staining of SOX17 in the endodermal cells, NCAM in the mesodermal cells, and PAX6 in the ectodermal cells. Scale bars, 400 μm.
